# Nuclear envelope assembly relies on CHMP-7 in the absence of BAF-LEM-mediated hole closure

**DOI:** 10.1101/2023.07.06.547980

**Authors:** Sarah R. Barger, Lauren Penfield, Shirin Bahmanyar

## Abstract

Barrier-to-autointegration factor (BAF) is a DNA binding protein that crosslinks chromatin to assemble the nuclear envelope (NE) after mitosis. BAF also binds the Lap2b-Emerin-Man1 (LEM) domain family of NE proteins to repair interphase ruptures. The NE adaptors to ESCRTs, LEMD2-CHMP7, seal NE holes surrounding mitotic spindle microtubules (MTs), but whether NE hole closure in mitosis involves BAF-LEM binding is not known. Here, we analyze NE sealing after meiosis II in *C. elegans* oocytes to show that BAF-LEM binding and LEM-2^LEMD2^-CHMP-7 have distinct roles in hole closure around spindle MTs. LEM-2/EMR-1^emerin^ function redundantly with BAF-1 to seal the NE. Compromising BAF-LEM binding revealed an additional role for EMR-1 in maintenance of the NE permeability barrier and an essential role for LEM-2-CHMP-7 in preventing NE assembly failure. The WH domain of LEM-2 recruits the majority of CHMP-7 to the NE in *C. elegans* and a LEM-2 -independent pool of CHMP-7, which is mostly enriched in the nucleoplasm, also contributes to NE stability. Thus, NE hole closure surrounding spindle MTs requires redundant mechanisms that safeguard against failure in NE assembly to support embryogenesis.

## Introduction

The nuclear envelope (NE) is a domain of the endoplasmic reticulum (ER) that serves as a mechanically stable semi-permeable barrier to the genome (Ungricht and Kutay, 2017). The outer and inner nuclear membrane of the NE encase a lumen that is shared with the ER. The inner nuclear membrane (INM) has a unique protein composition and is associated with a meshwork of filamentous nuclear lamins. Each cell division, the NE and nuclear lamins disassemble to release mitotic chromosomes for capture by spindle microtubules (MTs). After chromosome segregation, the NE forms from ER-derived membranes. Assembly of nuclear pore complexes (NPCs) and closure of NE holes establish the NE permeability barrier. Failure in the barrier function of the NE can lead to DNA damage and disrupt genome regulation, highlighting the importance of understanding mechanisms that seal the NE after mitosis (Gauthier and Comaills, 2021; Rodriguez-Muñoz et al., 2022).

NE formation relies on the dsDNA binding protein, Barrier to Autointegration Factor (BAF), that dimerizes to crosslink DNA and bind to a subset of integral INM proteins and lamins (Sears and Roux, 2020). Immediately after exit from mitosis, dephosphorylation of BAF promotes its high affinity association with segregated chromosome masses (Ahn et al., 2019; Asencio et al., 2012; Marcelot et al., 2021; Snyers et al., 2018). BAF dimers bridge DNA segments to ‘glue’ individual chromosomes together (Samwer et al., 2017). Integral membrane LAP2-Emerin-MAN (LEM)-domain proteins bind to a groove at the BAF dimer junction through a conserved 40 amino acid ‘LEM-domain’ to tether associated ER membranes around the chromatin surface (Barton et al., 2015; Cai et al., 2007; Lin et al., 2000). Nascent nuclear membranes first wrap the exposed region of the segregated chromatin mass that is unoccupied by spindle microtubules (called the ‘non-core’ domain) (Liu and Pellman, 2020). The majority of NPCs assemble in the non-core domain to initiate nuclear transport.

After the initial phase of NE formation, BAF accumulates at the ‘core’ domain of the chromatin mass, which is the densely occupied by spindle MTs (Haraguchi et al., 2008). BAF recruits and concentrates LEM-domain proteins, including LEMD2 and emerin, to the core domain where NE sealing of holes occurs (Haraguchi et al., 2008; von Appen et al., 2020). The LEM-domain protein LEMD2 contributes to NE sealing through its C-terminal winged helix (WH) domain that directly binds and activates the conserved ESCRT-II/ESCRT-III hybrid protein CHMP7 (Gatta et al., 2021; Gu et al., 2017; von Appen et al., 2020). The WH of LEMD2 copolymerizes with CHMP7 to form 50-100 nm rings *in vitro* and it is thought that their assembly on the cytosolic surface of NE holes restricts diffusion of macromolecules (Gatta et al., 2021; von Appen et al., 2020). CHMP7 serves as the NE adaptor for downstream ESCRT-III membrane remodeling machinery including the spiral filament protein CHMP4B/VPS32 and the microtubule severing protein Spastin that coordinate spindle disassembly with fusion of small (<100 nm) NE holes (Vietri et al., 2015; Ventimiglia et al., 2018; von Appen et al., 2020). Functions for other LEM-domain proteins at the core domain are less well understood.

The essential role for BAF in NE assembly as well as the functional redundancy of multiple LEM-domain proteins have made it challenging to test if BAF binding to LEM proteins serves a function aside from downstream ESCRT recruitment. In mitotically dividing cells, expression of a dimerization mutant in BAF, deficient in both DNA crosslinking and LEM-domain binding, results in hyper-micronucleation where membranes wrap individual chromosomes because they are not crosslinked (Samwer et al., 2017). Expression of a BAF mutant (BAF-L58R) that selectively prevents binding to LEM proteins did not cause micronucleation, demonstrating that BAF-LEM binding is not essential for formation of a single nucleus; however, whether sealing during NE formation is impaired in the absence of BAF-LEM binding was not tested. Expression of BAF-L58R does not support repair of large holes that result from ruptures indicating that BAF-LEM binding mediates NE sealing in interphase cells (Young et al., 2020). Importantly, CHMP7 accumulates at nuclear rupture sites but is not required for repair of ruptures (Halfmann et al., 2019). Thus, in interphase, BAF-LEM binding has a distinct role from CHMP7 in NE sealing, particularly of large NE holes that do not contain MTs (Halfmann et al., 2019; Young et al., 2020). Whether BAF-LEM binding and CHMP7 have separate roles in sealing NE holes that surround spindle MTs in mitosis is not known. Interestingly, LEMD2 binds MTs directly, which contributes to its enrichment at the core domain for NE sealing in mitosis (von Appen et al., 2020), so there may be a separable function for LEMD2-CHMP7 from BAF-LEM binding that depends on the presence of MTs.

Here, we take advantage of the opportunities provided by *C. elegans* to combine genetics and live imaging to dissect the contributions of BAF to NE sealing and the relationship of sealing to successful nuclear assembly. We focus on NE assembly after anaphase of meiosis II in fertilized oocytes to determine the shared and unique functions for BAF, LEM-domain proteins and CHMP7 in sealing of the large hole that surrounds spindle MTs, which establishes the permeability barrier in the first instance of NE formation for the developing embryo (Penfield et al., 2020).

Our previous work defined the assembly dynamics of *C. elegans* LEM-2 (human LEMD2) and presence of CHMP-7 (human CHMP7) at the micron-scale sized hole that surrounds the asymmetric meiotic spindle (Penfield et al., 2020). Importantly, closure of the post-meiotic NE hole does not require CHMP-7 (Penfield et al., 2020), but whether it requires the accumulation of membrane-bound LEM-domain proteins was not known. The regulation and requirement for BAF (Ce BAF-1) in nuclear assembly is conserved in *C. elegans* (Gorjanacz et al., 2007; Margalit et al., 2005). Furthermore, in contrast to human cells that contain seven LEM-domain proteins, *C. elegans* contain only two integral membrane LEM-domain proteins, EMR-1 (human emerin) and LEM-2, as well as a non-transmembrane containing LEM-domain protein LEM-3 (human ANKLE1), none of which are essential genes (Barton et al., 2015; Lee et al., 2000). Loss of both *lem-2* and *emr-1* results in embryonic lethality, which suggested that BAF-1 mediates its essential functions in nuclear assembly through redundant recruitment of these LEM-domain proteins (Liu et al., 2003).

We introduced the conserved BAF-L58R separation-of-function mutation at the endogenous locus of *C. elegans baf-1* to disrupt BAF-LEM binding. This mutant background allowed us to analyze the contribution of BAF-LEM binding to NE sealing and assembly. Our work reveals that EMR-1 and LEM-2 function redundantly in binding to BAF-1 to facilitate NE sealing around spindle MTs. We demonstrate that LEM-2-CHMP-7 are critical to stabilize the NE and safeguard against failure in NE assembly when BAF-LEM binding is compromised. Our genetic studies further reveal unique functions for LEM-2 and EMR-1 in maintenance of the NE permeability barrier and NE assembly. Thus, our work reveals redundant and distinct roles for multiple key players in NE formation and dissects the relationship between NE sealing and NE stability to support early embryonic development.

## Results

### BAF-1 dynamics and regulation after chromosome segregation in meiosis II and mitosis

Fertilization by haploid sperm triggers two rounds of meiosis in prophase I arrested *C. elegans* oocytes to produce the haploid pronucleus as well as two extruded polar bodies (Fig. 1A, left; (Fabritius et al., 2011)). The haploid oocyte-derived pronucleus forms as the acentriolar spindle elongates between segregated chromosomes after anaphase II (Fig. 1A, right). The NE initially assembles on the side of chromatin that is farthest from the acentriolar meiotic spindle and wraps around chromatin to form a sealing plaque akin to the ‘core domain’ that surrounds persisting spindle microtubules in mitosis (Fig. 1A, 100s; (Penfield et al., 2020)). The plaque condenses as the spindle dissipates (Fig. 1A, 200s) and then disperses as the pronucleus rapidly expands (Fig. 1A, 400s). In contrast to the oocyte-derived pronucleus, the haploid sperm-derived pronucleus, at the opposite end of the embryo, does not undergo closure of a large hole around spindle microtubules (Fig. 1A, left). Thus, the fertilized *C. elegans* zygote provides the opportunity to directly compare nuclear assembly with and without closure of a large hole around spindle microtubules in a shared cytoplasm (Fig 1A; (Penfield et al., 2020)). Furthermore, analyzing the first instance of NE formation allows us to eliminate confounding effects of prior rounds of failed NE assembly. Once formed, the oocyte-derived and sperm-derived pronuclei meet at pseudocleavage (PC) regression and progress to the first mitotic division (Fig. 1A; (Oegema, 2006)).

**Figure 1.**
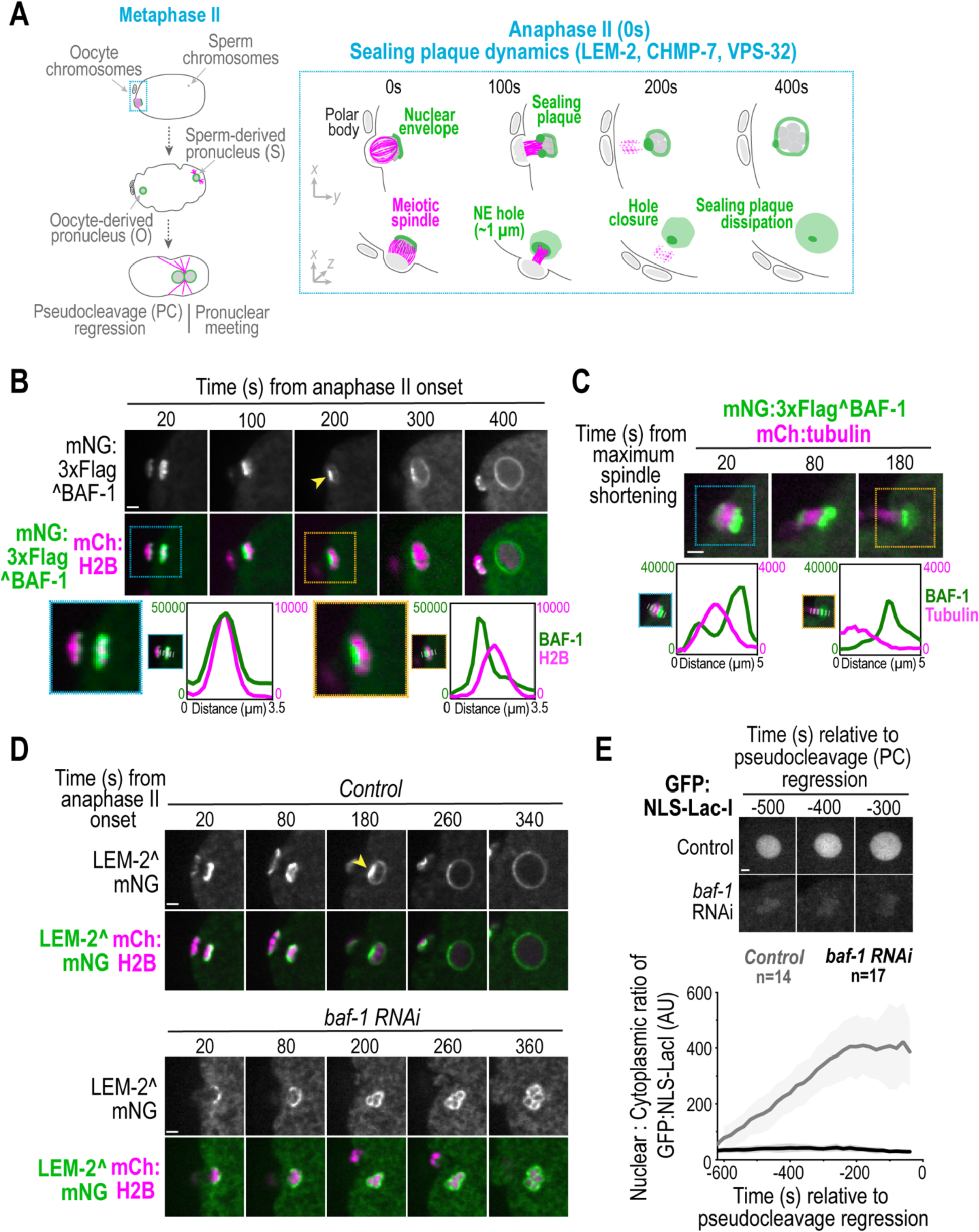
BAF-1 is required for sealing plaque formation during meiosis II in *C. elegans*. (A) Left, Schematic representation of oocyte- and sperm-derived pronuclear formation and migration after meiosis II. Right (box), schematic of sealing plaque dynamics relative to anaphase II. (B) Spinning disk confocal images from time lapse series of mNG^BAF-1 dynamics relative to anaphase II. Yellow arrowhead marks sealing plaque. Below, zoom insets of select frames for background-corrected line scans of indicated markers at indicated timepoints. Time in seconds relative to anaphase II onset. (C) Spinning disk confocal images from time lapse series of mNG^BAF-1 (green) and meiotic spindle microtubules (magenta). Below, zoom insets of select frames for background-corrected line scans of indicated markers at indicated timepoints. Time in seconds relative to maximum spindle shortening. (D) Spinning disk confocal images from time lapse series of LEM-2^mNG in indicated conditions. Yellow arrowhead marks sealing plaque. Time in seconds relative to anaphase onset. (E) Above, Spinning disk confocal images of GFP:NLS:LacI fluorescence in oocyte-derived pronucleus in indicated conditions. Below, plot (average ± SD) of normalized nuclear GFP:NLS-LacI fluorescence for indicated conditions. Time in seconds relative to pseudocleavage (PC) regression. n = # of embryos. Scale bars, 2 μm.

We generated a strain of *baf-1* tagged at its endogenous locus with mNeonGreen (mNG) and 3XFlag at the N-terminus (‘mNG^BAF-1’) to monitor BAF-1 dynamics at the sealing plaque. Embryo production and viability were unaffected in this strain suggesting that the fusion protein does not significantly interfere with BAF-1 function (Fig S1A). mNG^BAF-1 localizes on oocyte chromatin at anaphase II onset and transitions to a bright focus at the sealing plaque as the adjacent meiotic spindle lengthens and dissipates (Fig 1B, C). mNG^BAF-1 at the sealing plaque localized uniformly at the nuclear rim ∼400s following anaphase II onset (Fig 1B). mNG^BAF-1 associated with sperm chromatin transitioned to a small focus that enriched along the nuclear rim with similar dynamics as the oocyte-derived pronucleus (Fig. S1B; Movie 1). Thus, universal signaling mechanisms in the shared cytoplasm of the fertilized oocyte triggered by anaphase II control BAF-1 dynamics on chromatin and the nuclear envelope. The LEM-4-like protein (*lem-4*, human ANKLE-2) is an adaptor for PP2A that functions to dephosphorylate BAF-1, which enhances its chromatin-association immediately after mitotic exit (Asencio et al., 2012; Snyers et al., 2018; Marcelot et al., 2021) (Fig. S1C). Reducing *lem-4* by RNAi-depletion to maintain BAF-1 in a phosphorylated state reduced mNG^BAF-1 accumulation at segregated meiotic chromosomes (Fig. S1D) and delayed its accumulation on segregated mitotic chromosomes, as previously reported (Fig S1E; (Asencio et al., 2012). Thus, LEM-4 likely regulates BAF-1 dynamics in meiosis II similar to mitotically dividing cells.

### Enrichment of LEM-2 and EMR-1 at the sealing plaque in meiosis II requires BAF-1

We tested whether BAF-1 controls the dynamics of LEM-2 and EMR-1 (emerin) at the reforming NE and sealing plaque. We generated strains with *lem-2* and *emr-1* endogenously tagged with mNG using CRISPR/Cas9 editing. LEM-2^mNG and mNG^EMR-1 dynamics to the nuclear rim and enrichment at the sealing plaque were similar to mNG^BAF-1 and to prior reports for a LEM-2 transgene (Fig 1D and S1F; (Penfield et al., 2020)). However, they did not accumulate at a sealing plaque in *baf-1* RNAi-depleted embryos and instead membranes surrounded individual chromosomes that appeared condensed resulting in a multi-lobed oocyte pronucleus (Fig 1D and S1F, Movie 2). Furthermore, the initial levels of LEM-2-occupied membranes on chromatin during anaphase-II onset were significantly reduced in *baf-1* RNAi-depleted embryos (Fig. 1D and S1G). The substantial ER-associated pool of mNG^EMR-1 made comparable measurements unreliable (Fig. S1F). Embryos depleted of *baf-1* did not accumulate GFP:Nuclear localization signal (NLS)-LacI in the nucleus, indicating a failure in NE sealing (Fig. 1E). Thus, similar to penetrant RNAi-depletion of BAF in mammalian cells exiting mitosis (Samwer et al., 2017), BAF-1 is necessary to form a single nucleus after meiosis II in *C. elegans.* Additionally, BAF-1 directs formation of the sealing plaque containing LEM-2 and EMR-1, which prompted us to use genetic analysis to understand how BAF-1’s recruitment of LEM-domain proteins contributes to sealing of the large post-meiotic NE hole.

### A mutation in BAF-1 that prevents LEM-domain binding reduces its NE localization and is non-essential in *C. elegans*

To understand the contribution of BAF-LEM binding in closure of the large hole that surrounds meiotic spindle microtubules, we introduced mutations in conserved amino acid residues in *C. elegans baf-1* that have been shown to serve as separation-of-function mutations in human BAF (Samwer et al., 2017; Halfmann et al., 2019; Young et al., 2020; Fig 2A). BAF-G47E disrupts BAF dimerization and binding to LEM-domain proteins while BAF-L58R selectively inhibits LEM-domain binding (Fig. 2A, B; (Halfmann et al., 2019; Samwer et al., 2017; Young et al., 2020)).

**Figure 2.**
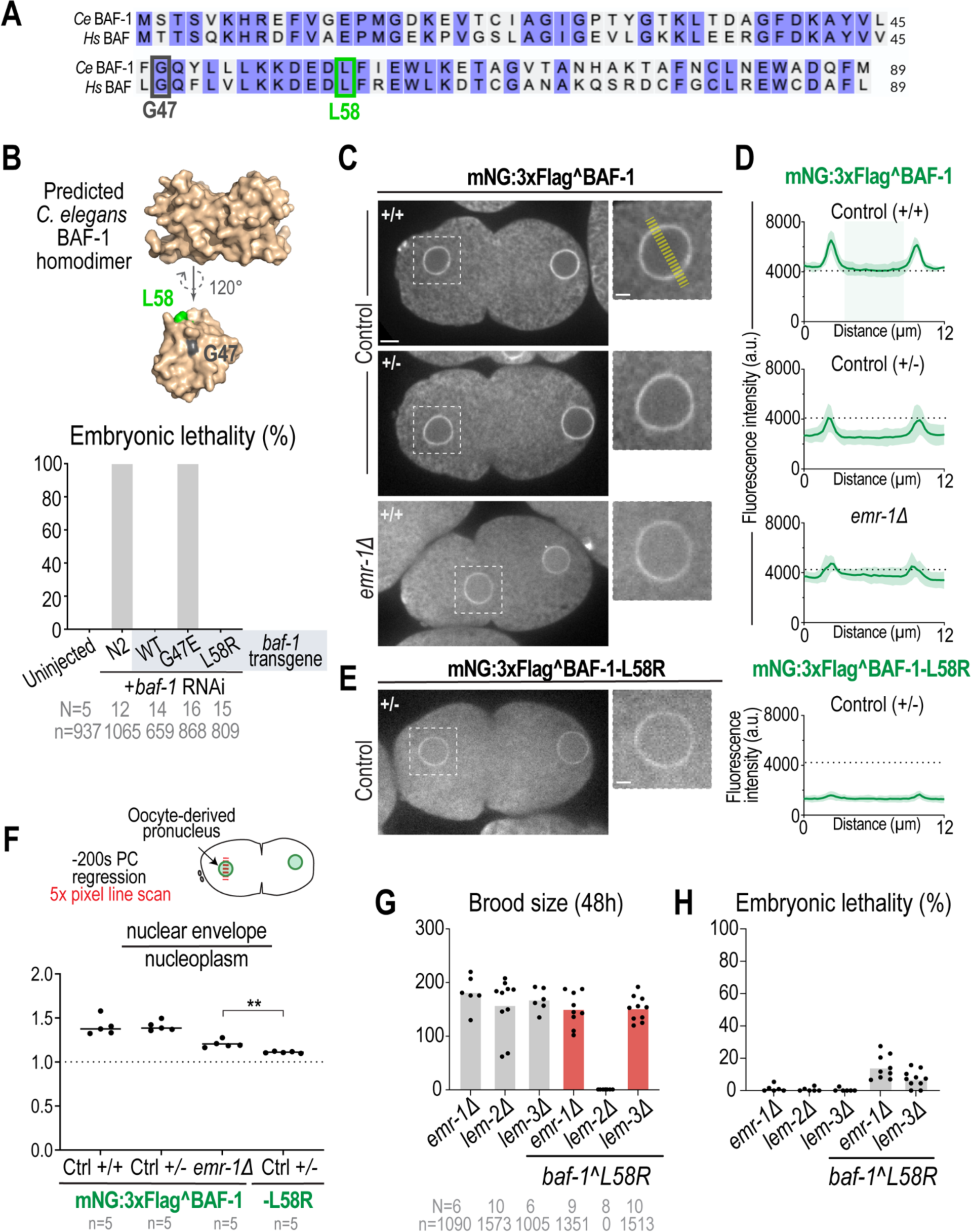
A mutation in BAF-1 that selectively inhibits LEM-domain binding is non-essential in *C. elegans.* (A) Amino acid sequence alignment of human BAF and *C. elegans* BAF-1. Identical amino acid residues shaded in purple; mutations in indicated amino acid residues are boxed. (B) Above, 3D space filling model of predicted *C. elegans* BAF-1 homodimer generated using (Miradita et al., 2022; Goddart et al., 2018; Pettersen et al., 2021) and monomer using AlphaFold Protein Structure Database (Jumper et al., 2021; Varadi et al., 2022). Monomer rotated 120 degrees on the x-axis relative to dimer with amino acid residues mutated in this study highlighted. Below, Plot of percentage of embryonic lethality in indicated conditions. N = # of worms, n = # of embryos. (C-E) Left, Spinning disk confocal images of mNG^BAF-1 at 200 sec prior to pseudocleavage (PC) regression in indicated conditions. Homozygous (+/+) or heterozygous (+/-) insertion of mNG at endogenous *baf-1* locus marked in images. Right, Plots of background-corrected line scan (average ± SD) of mNG^BAF-1 (n = 5 embryos). Fluorescence signal from nucleoplasm shaded in green in control. Scale bars, 5 μm; Scale bar in zoom insets, 2 μm. (F) Above, Schematic representation of line scan analysis. Below, Plot representing ratio of mNG signal at nuclear envelope (average of peak signals) and nucleoplasm (average signal between peaks) from line scan analysis in indicated conditions. Statistical significance determined by Mann-Whitney test (**=p-0.0079). n = # of embryos. (G, H) Plot representing embryonic lethality (G) and brood size (H) in indicated conditions. N = # of worms, n = # of embryos.

We first introduced the mutations into RNAi-resistant *baf-1* transgenes to avoid embryonic lethality and sterility that could result from significant disruption of *baf-1* function (Fig. S2A). RNAi-depletion of *baf-1* caused 100% embryonic lethality (Fig. 2B), as expected (Gorjanacz et al., 2007; Margalit et al., 2005; Zheng et al., 2000), and this was rescued in the presence of the WT re-encoded *baf-1* transgene, but not the dimerization deficient *baf-1(G47E)* mutant transgene (Fig. 2B). The *baf-1(L58R*) mutant re-encoded transgene supported viability in *baf-1* RNAi-depleted embryos (Fig. 2B). We therefore introduced the *baf-1(L58R)* mutation at the endogenous locus (Fig. S2B). The *baf-1(L58R)* mutant worm expressed normal levels of BAF-1 protein (Fig. S2C), did not cause embryonic lethality, and resulted in a slight reduction in brood size (Fig. S2D). Together these data indicate that BAF-LEM binding is not essential for germline development or embryogenesis.

We next tested if BAF-LEM binding is required for BAF-1 localization at the INM. BAF localizes to both the nucleoplasm and INM; LEM-domain proteins bind BAF at the INM (Haraguchi et al., 2007; Liu et al., 2003), so we expected to observe reduced BAF-1-L58R mutant protein at the NE, but not the nucleoplasm (Halfmann et al., 2019; Samwer et al., 2017; Young et al., 2020). In the 1-cell stage embryo, mNG^BAF-1 is enriched at the NE of the oocyte- and sperm-derived pronuclei, and localizes to the nucleoplasm, ER, and diffusely in the cytoplasm (Fig. 2C). Our attempts to homozygous a *baf-1(L58R)* mutant animal tagged at the endogenous locus with mNG were unsuccessful suggesting that the mNG tag together with the L58R mutation compromise BAF-1 function at the organismal level. To bypass this issue, we quantified the NE and nucleoplasmic fluorescence signal of heterozygous embryos carrying one copy of mNG tagged *baf-1* and one untagged copy of wild type *baf-1* (Fig. 2C-F). In heterozygous wild type mNG^ *baf-1* / *baf-1* embryos, the NE and nucleoplasmic mNG fluorescence signal was approximately half that of the homozygous mNG tagged wild type *baf-1* strain (Fig. 2C, D). The NE to nucleoplasmic ratio of the mNG fluorescence was the same in both homozygous and heterozygous strains, so even with half the amount of protein the proportion of wild type BAF-1 at the NE versus the nucleoplasm remains constant (Fig. 2F).

Both the NE and nucleoplasmic mNG fluorescence levels were significantly lower in heterozygote mNG^*baf-1*(L58R)/ *baf-1* than in wild type mNG^*baf-1*/*baf-1* embryos (Fig. 2E), although protein levels of untagged BAF-1-L58R in the *baf-1(L58R)* mutant strain that we use throughout this study are unchanged (Fig. S2C). The lower NE:nucleoplasmic ratio reflected a greater decrease of the mNG^BAF-1-L58R mutant protein at the NE than nucleoplasm (Fig. 2E, F). The faint mNG^BAF-1-L58R localization at the nuclear rim may be through dimerization with untagged wild type BAF-1 or through association with the nuclear lamina or an unidentified NE adaptor (Holaska et al., 2003; Kono et al., 2022; Samson et al., 2018). ER-localized signal was also lost in mNG^ *baf-1*(L58R)/ *baf-1* heterozygote embryos indicating that BAF-1 associates with the ER through binding to LEM domain proteins (Fig. 2E). Reduced nuclear rim, but unchanged nucleoplasmic signal, of mNG^BAF-1 in *emr-1Δ* embryos, as reflected in the lower NE:nucleoplasmic ratio (Fig. 2C, D), is consistent with past findings that emerin contributes to BAF-1 binding at the INM (Asencio et al., 2012). We were unable to assess the localization of mNG^BAF-1-L58R upon loss of *lem-2* because of the synergistic sterility phenotype that results from this genetic background (see below Fig. 2G). Together, we conclude that the *C. elegans* BAF-1-L58R mutant is compromised in its NE-association, as previously shown for the mutant human BAF homologue (Halfmann et al., 2019; Samwer et al., 2017; Young et al., 2020), and that multiple LEM-domain proteins recruit BAF-1 to the INM.

### Distinct functions for LEM proteins in embryo and germline development revealed by the *baf-1(L58R)* mutant

We crossed the *baf-1(L58R)* strain to deletion alleles in each of the three *C. elegans* LEM-domain genes (*emr-1, lem-2,* and *lem-3*) because we predicted that if LEM proteins have redundant or related functions to BAF-LEM binding, then a double mutant (e.g. *emr-1Δ; baf-1(L58R))* would result in synergistic phenotypes (e.g. increased lethality). Alternatively, if LEM proteins function only through binding to BAF-1, then a double mutant with *baf-1(L58R)* would not cause a significant increase in embryo viability or production. Deletion of *emr-1* or a *lem-3* mutant allele with reduced function in the *baf-1(L58R)* mutant strain resulted in only a slight decrease in brood size and embryonic viability (Fig. 2G, H) suggesting that these LEM proteins do not have a function that is redundant with BAF-LEM binding to support embryo and germline development. In contrast, deletion of the *lem-2* gene locus did not cause lethality or sterility on its own (Fig. 2G, H) but the *lem-2Δ*; *baf-1(L58R)* double mutant inhibited germline development resulting in sterility in 100% of worms (Fig. 2G, S2I). Note that we found an annotation error in the *lem-2(tm1582)* mutant allele, which has been widely used and interpreted as a *lem-2* deletion strain (Barkan et al., 2012; Bone et al., 2014; Dittrich et al., 2012; González-Aguilera et al., 2014; Harr et al., 2020; Morales-Martínez et al., 2015; Penfield et al., 2020; Shankar et al., 2022), in which the 3’ region that encodes for the WH domain is intact (Fig. S2F) (Davis et al., 2022). We therefore generated a CRISPR-Cas9 gene edited strain that deletes the entire *lem-2* gene locus for this study (Fig. S2G, H). Together, these data indicate that LEM-2, but not EMR-1 or LEM-3, has a redundant function to BAF-LEM binding in germline development, and further verify that the BAF-1-L58R mutant protein is functionally compromised.

### Abnormal dynamics of endogenously tagged LEM-2 and EMR-1 at the reforming NE and sealing plaque in *baf-1(L58R)* mutants

We next tested whether compromising BAF-LEM protein interactions impacted the recruitment of endogenous LEM-2^mNG and EMR-1^mNG to the reforming post-meiotic NE (Fig. 3, Fig S3A). LEM-2^mNG appeared on the nascent NE at anaphase II onset at lower levels in *baf-1(L58R)* mutant embryos compared to control embryos (Fig. 3A, B, Movie 3), similar to *baf-1* depletion (Fig. S1G). Instead of the organized sealing plaque that forms directly adjacent to the meiotic II spindle under control conditions (100s, Fig. 3A), LEM-2^mNG formed smaller foci around condensed chromatin that coalesced into a single punctum with lower fluorescence intensity levels and delayed appearance in *baf-1(L58R)* mutants compared to controls (Fig. 3C-D). Furthermore, the punctum of LEM-2^mNG in *baf-1(L58R)* mutant oocytes persisted ∼ 300s longer than the sealing plaque in control embryos (Fig. 3E). Fully formed oocyte-derived pronuclei in *baf-1(L58R)* mutant contained lower levels of LEM-2^mNG at the NE (Fig. S3B, C) despite equal levels of global LEM-2 protein (Fig S3D). Similar observations were made for mNG^EMR-1 dynamics in *baf-1(L58R)* mutants (Fig. 3F, Fig. S3B, C). Additionally, compromised BAF-LEM binding resulted in delayed LEM-2^mNG on nascent nuclear membranes following the first mitotic division that did not fully enrich at the ‘core’ domain (Fig. S3E-G). Together, these data confirm that BAF-1 recruits and organizes LEM proteins to the sealing plaque through its BAF-LEM domain binding function. The fact that LEM-2 and EMR-1 can localize to the NE independent of BAF-1 binding, albeit at significantly lower levels (Fig S3B, C), supported the possibility that these proteins have functions independent of BAF-1-mediated recruitment at the NE.

**Figure 3.**
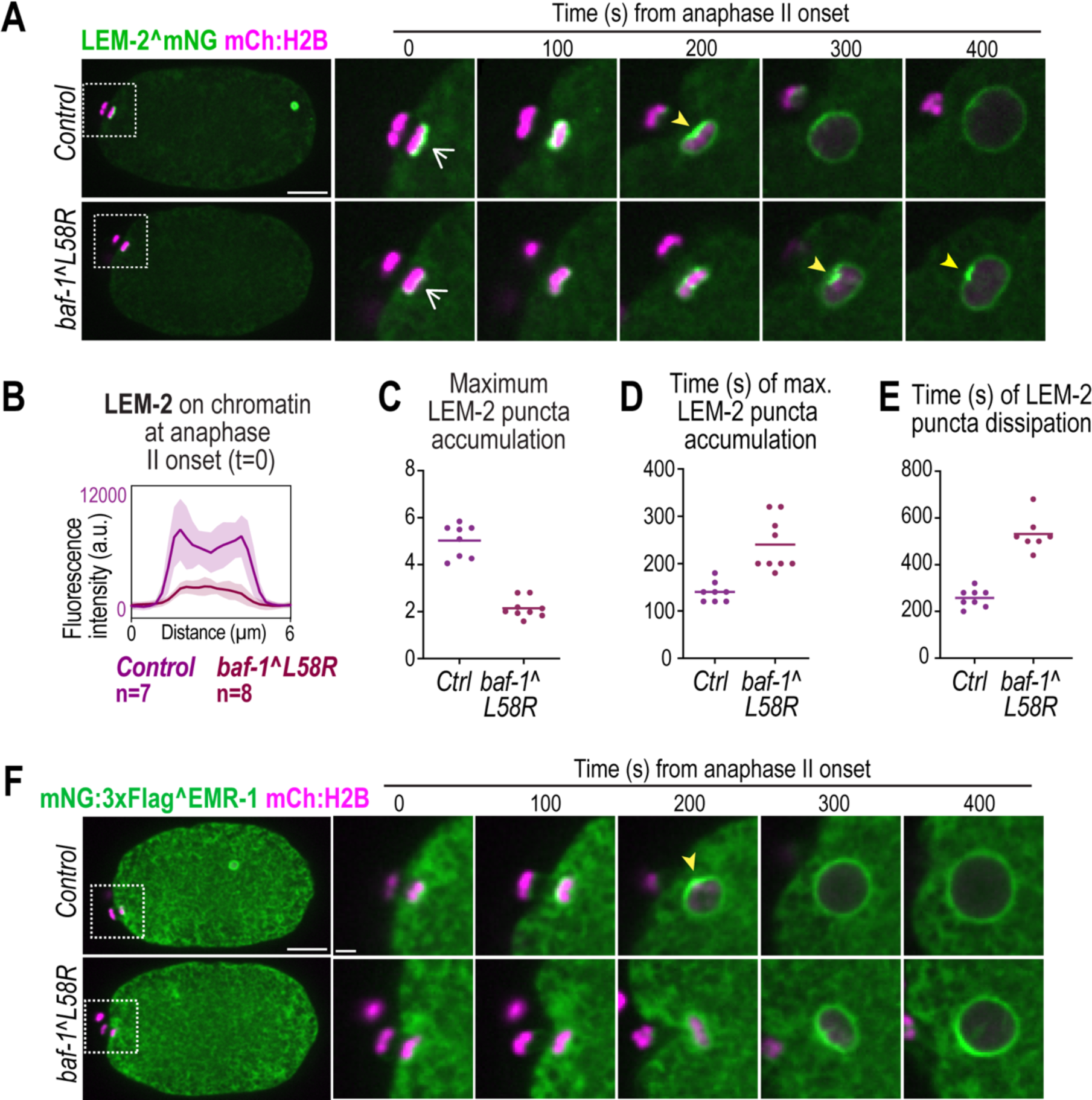
BAF-LEM interactions orchestrate organized and timely recruitment of sealing proteins during nuclear envelope formation. (A) Spinning disk confocal images from time lapse series of LEM-2^mNG during oocyte pronuclear formation in indicated conditions. (B) Plot of background-corrected line scan analysis (average ± SD) of LEM-2^mNG on at nuclear rim at anaphase II onset (white arrows at t = 0s) in indicated conditions. n = # of embryos. (C-D) Plots (average + replicates) representing maximum LEM-2 fluorescence signal at nuclear rim from time series in (C), time in seconds relative to anaphase onset II of maximum LEM-2^mNG fluorescence signal in (C) shown in (D), and time in seconds relative to anaphase onset II of LEM-2^mNG puncta dissipation shown in (E). (F) Spinning disk confocal images from time lapse series of mNG^EMR-1 in indicated conditions. Yellow arrows point to enrichment sites. Scale bars of whole embryo image, 10 μm, zoom insets scale bar, 2 μm.

### LEM-2 and EMR-1 compensate for each other in binding to BAF-1 to seal the NE around spindle MTs

We compared NE assembly and sealing in the oocyte versus sperm-derived pronuclei to determine if there is a unique requirement for BAF-LEM binding in closure of the large NE hole surrounding spindle MTs (Fig. 4). Live imaging of a general ER marker (SP12:GFP) and chromosomes (mCh:Histone(H)2B) showed that 100% of both oocyte and sperm-derived pronuclei RNAi-depleted of *baf-1* are multi-lobed (Fig. 4B), while in *baf-1(L58R)* mutants a proportion of oocyte-but not sperm-derived pronuclei were malformed and had faint intranuclear membranes (Fig. 4A, B). Reducing *lem-4* levels to prevent dephosphorylation of BAF-1 mildly impacted nuclear shape on its own (Fig. 4B), but resembled a penetrant BAF-1 depletion in *baf-1(L58R)* mutants, which was specific to the oocyte-derived pronucleus (Fig. 4A, B, Movie 4). These data indicate that there is a greater reliance on BAF-LEM binding for nuclear formation when a large hole that surrounds spindle microtubules must be sealed. Nuclear formation in mitotic *baf-1-L58R* embryos was delayed and resulted in smaller nuclei (Fig. S4A, B) suggesting that BAF-LEM binding contributes to NE assembly in both meiosis and mitosis in *C. elegans*.

**Figure 4.**
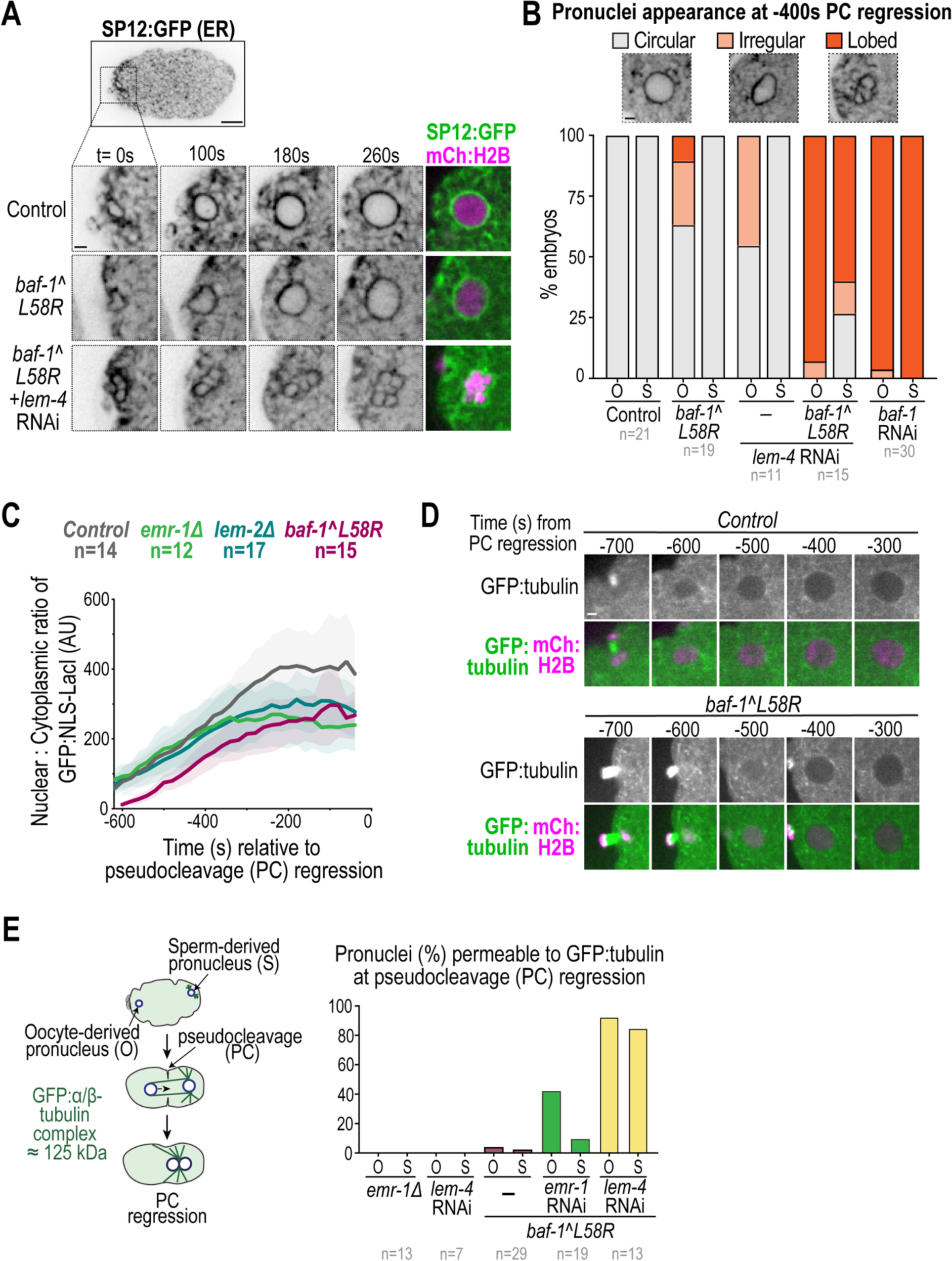
Reliance on BAF-LEM binding to seal the NE large hole surrounding meiotic spindle microtubules. (A) Spinning disk confocal images from time lapse series of oocyte pronuclear formation marked by SP12:GFP (ER marker) and mCh:Histone(H)2B in indicated conditions. (B) Plot representing percentage of oocyte-derived (O) and sperm-derived (S) pronuclei categorized as circular, irregular, or lobed at −400 sec relative to pseudocleavage (PC) regression in indicated conditions. n = # of embryos. (C) Plot representing average ± SD of ratio of normalized GFP::NLS-LacI fluorescence in oocyte pronucleus for indicated conditions relative to PC regression. n = # of embryos. (D) Spinning disk confocal images from time-lapse series of nuclear GFP:α-tubulin during oocyte pronuclear formation in indicated conditions. Time in seconds relative to PC regression. (E) Left, schematic represents nulear exclusion of GFP:α-tubulin from pronuclei during pronuclear migration and pseudocleavage (PC) formation and regression. Right, plot representing percentage of oocyte-derived (O) and sperm-derived (S) pronuclei permeable to GFP:α-tubulin at pseudocleavage regression. n = # of embryos. Scale bars, 2 μm.

We next monitored the time course of nuclear import of a GFP:NLS reporter following anaphase II onset to PC regression in both the oocyte and sperm-derived pronuclei in *baf-1(L58R)* mutant embryos (Fig. 4C, S4C). The nuclear to cytoplasmic ratio of the GFP:NLS reporter directly following nuclear formation in the oocyte-derived, but not in the sperm-derived pronucleus, was significantly lower in *baf-1(L58R)* embryos compared to wild type embryos (Fig 4C, Fig. S4C). Furthermore, nuclear exclusion of GFP:α tubulin, which is actively exported from pronuclei (Hayashi et al., 2012) and serves as an indicator of closure of large NE holes (Penfield et al., 2020), was delayed, but not inhibited, in the *baf-1(L58R)* mutant oocyte-derived pronucleus (Fig 4D, E). Together, these data indicate that the timely closure of the large NE hole surrounding meiotic spindle MTs requires the ability of BAF to bind LEM proteins.

Deletion of *lem-2* did not impact the initial rate of nuclear import (Fig. 4C) nor nuclear assembly (Fig. S4D, E) following anaphase II, although the nuclear GFP:NLS signal did not reach the maximum wild type levels in these mutant embryos suggesting that LEM-2 may have a parallel function that in addition to binding to BAF-1 is required for maintenance of the NE permeability barrier. Sterility resulting from loss of *lem-2* in *baf-1(L58R)* worms prevented us from genetically testing the consequences on nuclear sealing resulting from loss of *lem-2* in *baf-1(L58R)* embryos.

Deletion of *emr-1* resulted in lower retention of nuclear GFP:NLS (Fig. S4D, E; Fig. 4C), similar to loss of *lem-2*. In *baf-1-L58R* pronuclei, RNAi-depletion of *emr-1* resulted in ∼42% of oocyte-derived containing nuclear GFP:α-tubulin at PC regression (Fig. 4E), suggesting that EMR-1 has an additional function to BAF-1 in stabilizing the NE barrier when NE sealing around spindle MTs is required. Both oocyte- and sperm-derived pronuclei were permeable to GFP:α-tubulin at PC regression upon RNAi-depletion of *lem-4* in the *baf-1(L58R)* mutant, further supporting the general requirement for both chromatin binding and LEM domain binding functions of BAF-1 to support stable nuclear assembly (Fig. 4A, B, E).

We conclude that LEM-2 and EMR-1 compensate for each other in binding to BAF-1 to ensure NE sealing around spindle MTs - loss of either gene alone results in milder defects in NE sealing than in the *baf-1(L58R)* mutant. Our data also reveal that there are unique functions for LEM-2 and EMR-1 that are distinct from BAF-LEM binding. EMR-1 has a parallel function to BAF-LEM binding required for maintenance of the NE barrier. The sterility phenotype that is specific to loss of *lem-2* in the *baf-1(*L58R) mutant indicates a unique function for LEM-2 from EMR-1 in germline development that is redundant to BAF-LEM binding.

### Nuclear envelope assembly around spindle MTs relies on CHMP-7 when BAF-LEM binding is compromised

The mild NE assembly defects and delayed but not failed sealing in the absence of BAF-LEM binding prompted us to test for other factors that may be functioning redundantly to prevent catastrophic loss of the NE permeability barrier. Our data showing that LEM-2 and EMR-1 are recruited to the NE even in the absence of BAF-LEM binding suggested that downstream ESCRT-mediated NE remodeling may support NE formation under these conditions. Furthermore, evidence in *C. elegans* suggests that CHMP-7 contributes to NE sealing and stability under conditions of increased lipid biogenesis or a weakened nuclear lamina (Penfield et al., 2020; Shankar et al., 2022). We did not detect defects in formation of the oocyte and sperm-derived pronuclei in embryos deleted for *chmp-7* (Fig. 5A, B), similar to our past results (Penfield et al., 2020). In contrast, in ∼35% of *chmp-711*; *baf-1(L58R)* double mutant embryos the oocyte-, but not sperm-, derived pronucleus appeared lobed (Fig. 5A, B and see also Fig. S5A). These NE assembly abnormalities led to ∼47% of *chmp-711*; *baf-1(L58R)* embryos containing collapsed oocyte-derived pronuclei at pronuclear meeting, whereas sperm-derived pronuclei appeared normal (Fig. S5A, Movie 5). Approximately half of *chmp-711*; *baf-1(L58R)* embryos do not survive to hatching (Fig. 5C), which is reflected in the severe mitotic defects observed in these mutants that appeared to compound with subsequent cell divisions (Fig. S5B). The enhanced phenotypes are specific to loss of *chmp-7* in the *baf-1(L58R)* mutant background in which both EMR-1 and LEM-2 binding to BAF-1 are compromised because *chmp-711*; *emr-111* and *chmp-711*; *lem-2Δ* mutant embryos are mostly viable (Fig. 5C) further indicating that binding of BAF-1 to EMR-1 and LEM-2 functions redundantly to promote hole closure.

**Figure 5.**
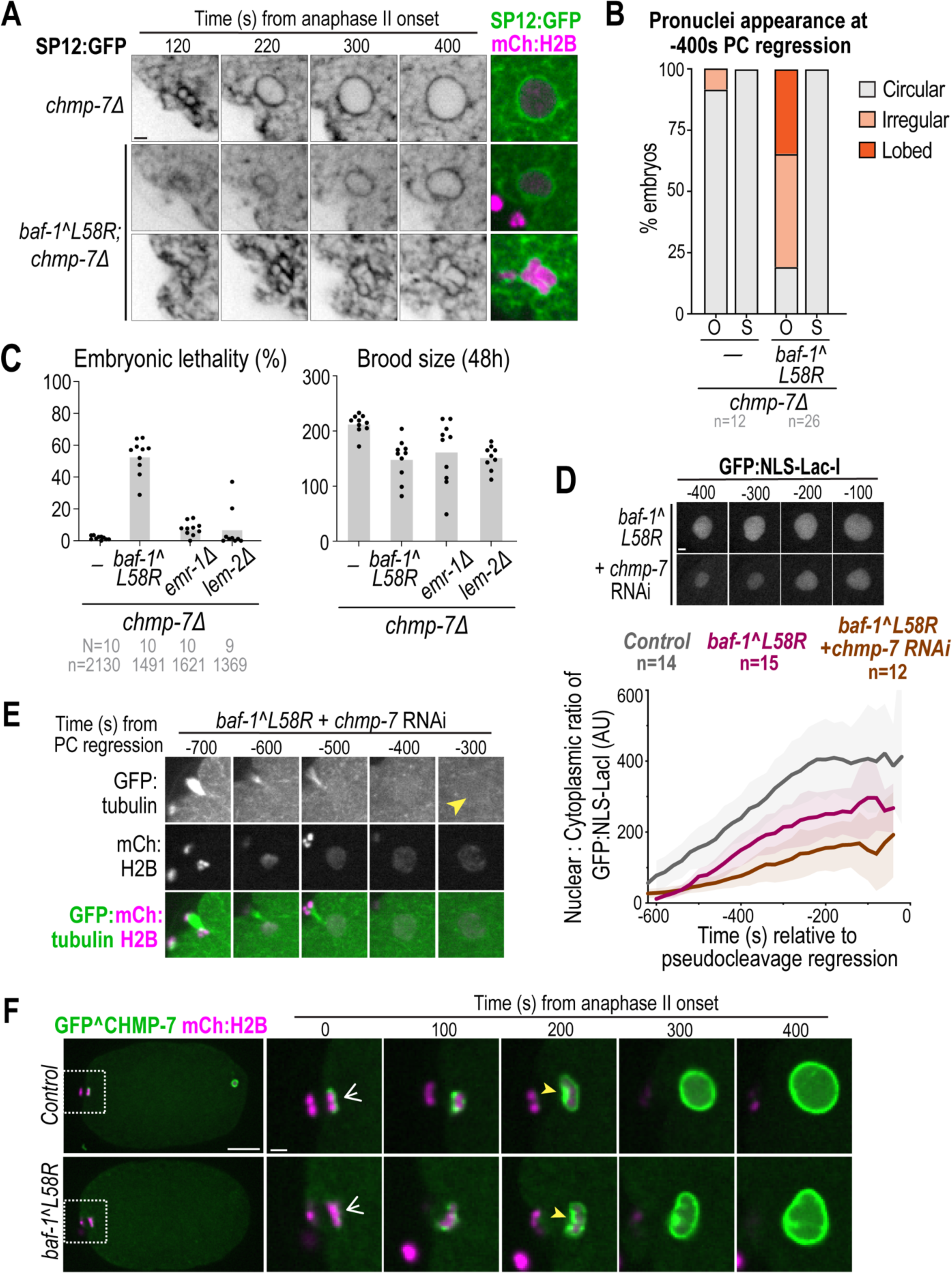
CHMP-7 maintains nuclear integrity when BAF-LEM-mediated hole closure is compromised. (A) Spinning disk confocal images from time lapse series of SP12:GFP (ER) and mCH:Histone(H)2B in indicated conditions during oocyte-derived pronuclear formation. (B) Plot representing percentage of oocyte-derived (O) and sperm-derived (S) pronuclei categorized as circular, irregular, or lobed at −400 sec relative to pseudocleavage (PC) regression in indicated conditions. n = # of embryos. (C) Plots representing average + replicates of percentage of embryonic lethality and brood size in indicated conditions. N = # of worms, n = # of embryos. (D) Plot representing average ± SD of normalized nuclear GFP:NLS-LacI fluorescence in indicated conditions. Time in seconds relative to pseudocleavage (PC) regression. Control and *baf-1-L58R* replicated from Fig. 4C. n = # of embryos. (E) Spinning disk confocal images from time-lapse series of GFP-α-tubulin in the oocyte-derive pronucleus in indicated conditions. Time in seconds relative to PC regression. Yellow arrow marks nuclear GFP-α-tubulin. (F) Spinning disk confocal images from time lapse series of mNG^CHMP-7 during oocyte pronuclear formation in indicated conditions. White arrows denote initial recruitment of GFP^CHMP-7 to chromatin and yellow arrows point to enrichment. Time in seconds relative to anaphase II onset. Scale bars of whole embryo images, 10 μm, all other scale bars, 2 μm.

While some *chmp-711*; *baf-1(L58R)* oocyte-derived pronuclei assembled into a normal shape (Fig. 5A, B, ∼19% “circular”), they did not establish or maintain nuclear accumulation of the GFP:NLS reporter (Fig. 5D). Furthermore, the majority (94%) of these pronuclei failed to exclude GFP:α-tubulin (Fig. 5E, S5C). Thus, CHMP-7 is required for successful, albeit delayed, establishment of the NE permeability barrier when BAF-LEM-domain binding is impaired. The fact that oocyte-derived pronuclei sometimes assemble under these conditions and ∼47% of embryos survive to hatching (Fig. 4C) suggests that the likelihood of NE assembly failure may depend on the extent of loss of the NE permeability barrier and stability of the NE hole. Together, these data show that CHMP-7 protects against failure in the NE barrier when NE hole closure around spindle MTs is defective.

### BAF-LEM binding controls the levels and dynamics of CHMP-7 during NE formation and sealing

To understand how CHMP-7 contributes to normal NE sealing and assembly in the absence of BAF-LEM binding, we first monitored the levels, dynamics and organization of endogenously tagged GFP^CHMP-7 at the reforming NE and sealing plaque after anaphase II onset. CHMP-7 is constitutively localized to the nuclear rim and in the nucleoplasm in *C. elegans* embryos (Shankar et al., 2022). In control embryos, nuclear rim-associated CHMP-7 appears during anaphase II onset, wraps around chromatin and enriches at the sealing plaque, similar to LEM-2 (Fig. 5F). GFP^CHMP-7 accumulates in the nucleoplasm ∼200s after anaphase II onset, concomitant with establishment of the nuclear permeability barrier (Fig. 5F, Movie 6). In the *baf-1(L58R)* mutant, GFP^CHMP-7 levels were initially lower at the nuclear rim (0s, Fig. S5D) and discrete CHMP-7 foci formed around condensed chromatin (Fig. 5F, 200 s), which coalesced into a disorganized focus rather than a distinct plaque (Fig. 5F). CHMP-7 also localized to the intranuclear membranes observed in assembled oocyte pronuclei of *baf-1(L58R)* mutant (400s, Fig. 5F). Lower levels and abnormal dynamics of GFP^CHMP-7 were also observed during NE formation in mitotic embryos of the *baf-1(L58R)* mutant (Fig. S5E, F), similar to LEM-2^mNG.

Thus, the localization and dynamics of endogenous CHMP-7 at the reforming NE after meiosis II and mitosis resembles that of LEM-2 and EMR-1 and is consistent with the role of BAF-1 in recruiting and organizing LEM-domain proteins that then regulate CHMP-7 localization and dynamics. However, the enhanced severity of *chmp-711*; *baf-1(L58R)* suggested that CHMP-7 functions independently of BAF-LEM binding to ensure NE sealing and assembly.

### LEM-2-dependent and independent pools of CHMP-7 contribute to post-meiotic NE assembly

CHMP-7 localizes to the NE in *baf-1(L58R)* mutants during reformation, albeit in a reduced and disorganized manner at initial timepoints, indicating recruitment of CHMP-7 to the NE is only partially dependent on BAF-LEM binding. Prior work had suggested that both LEM-2 and EMR-1 are redundantly required for CHMP-7 localization to the INM in *C. elegans* (Shankar et al., 2022); however, the misannotated *lem-2* mutant allele strain was used assess CHMP-7 dynamics (see Fig. S2F). We found that nuclear rim localization of GFP^CHMP-7 was not detectable in the CRISPR-Cas9 gene edited *lem-211* strain (Fig. 6A). GFP^CHMP-7 localization to the ER in mitotically dividing cells was also abolished in *lem-211* embryos (Fig. S6A). Deletion of the WH domain of *lem-2* using CRISPR-Cas9 gene editing further revealed that the WH of LEM-2 is responsible for recruiting CHMP-7 to the nuclear rim, similar to other systems (Fig. 6A) (Gu et al., 2017; von Appen et al., 2020). Deletion of *emr-1* in this genetic background did not result in a change in GFP^CHMP-7 localization (Fig 6A). Thus, LEM-2 is required to retain the majority of CHMP-7 at the INM in *C. elegans* embryos. We did observe a very slight nuclear rim fluorescence signal of GFP^CHMP-7 in *lem-2Δ* and *lem-2-ΔWH* mutants ∼160-180s after anaphase II onset in meiosis and anaphase onset in mitosis (Fig. 6B, S6C) indicating that some CHMP-7 can associate with the nuclear rim without LEM-2, which may be EMR-1-dependent. Neither LEM-2 nor EMR-1 are necessary for nuclear accumulation of CHMP-7 (Fig. 6A) and we observed increased nuclear GFP^CHMP-7 levels in *lem-2Δ* and *lem-2 11WH* oocyte-derived pronuclei (Fig. S6B). This suggested that the CHMP-7 pool that doesn’t localize to the nuclear rim in the absence of LEM-2 binding accumulates in the nucleoplasm. Together, these data demonstrate that the LEM-2 WH domain recruits CHMP-7 to the NE even in the absence of BAF-LEM binding, which may explain why CHMP-7 can compensate for defective NE sealing in the *baf-1(L58R)* background.

**Figure 6.**
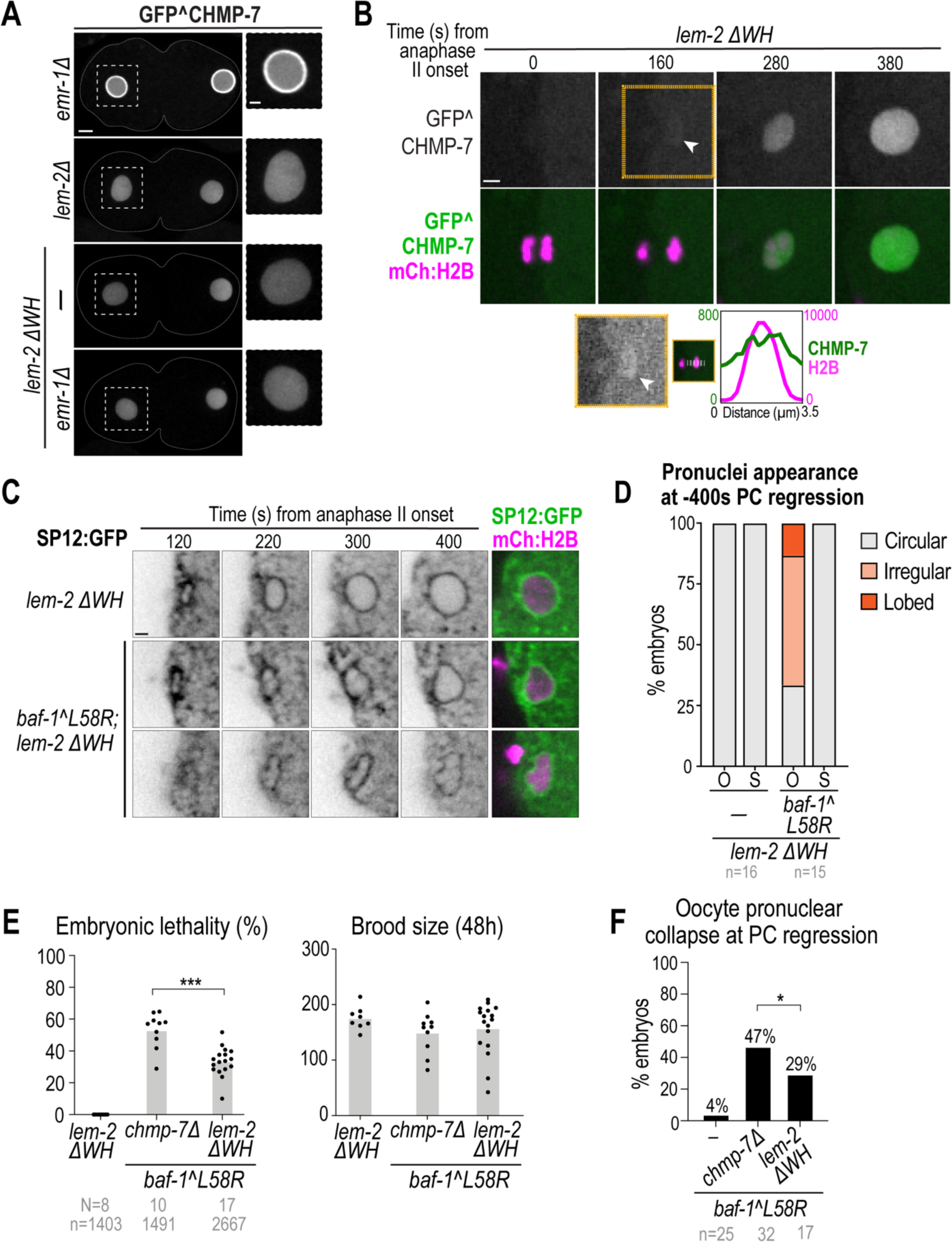
Compromising BAF-LEM binding reveals a role for CHMP-7 in nuclear stability independent of LEM-2 binding. (A) Left, Spinning disk confocal images of GFP^CHMP-7 in embryos at −200 sec relative to pseudocleavage (PC) regression in indicated conditions. Scale bars, 5 μm, zoom scale bars, 2 μm. (B) Spinning disk confocal images from time lapse series of GFP^CHMP-7 and mCH:Histone(H)2B during oocyte pronuclear formation in *lem-2 ΔWH* embryos. Time is in seconds relative to anaphase II onset. White arrow denotes faint membrane-association of GFP^CHMP7 prior to nuclear enrichment. Yellow outlined panel reproduced below montage with brightness/contrast adjusted. Plot represents background-corrected line scan of fluorescence of GFP^CHMP-7 and mCH:Histone(H)2B across reforming nucleus at 160 s post anaphase II onset. (C) Spinning disk confocal images from time lapse series of indicated markers in indicated conditions. (D) Plot representing percentage of oocyte-derived (O) and sperm-derived (S) pronuclei categorized as circular, irregular, or lobed at −400 sec relative to PC regression in indicated conditions. n = # of embryos. (E) Plots representing average + replicates of percentage of embryonic lethality and brood size in indicated conditions. N = # of worms, n = # of embryos. Statistical significance of indicated conditions determined by Welch’s t-test (***=p=0.0002). (F) Plot representing percentage of embryos that undergo oocyte pronuclear collapse in indicated conditions. n = # of embryos. Statistical significance of indicated conditions determined by Fisher’s exact test (*=p=0.0130).

We next tested whether the LEM-2 independent pool of CHMP-7 that is mostly nuclear-enriched can function to support nuclear assembly. Nuclear GFP^CHMP-7 accumulates during post-meiotic and mitotic NE formation, but fails to enrich at the sealing plaque or ‘core’ domain without LEM-2 (Fig. 6B, S6C, Movie 7). To test if the LEM-2-independent pools of CHMP-7 are functional in NE assembly, we monitored NE formation in *lem-2-ΔWH; baf-1(L58R)* double mutant embryos (Fig. 6C). A lower percentage of *lem-2-ΔWH; baf-1(L58R)* double mutant embryos displayed lobed oocyte-derived pronuclei (∼13%; Fig. 6C, D) compared to *chmp-711; baf-1(L58R)* double mutants (see Fig. 5B) suggesting the pool of CHMP-7 is partially functional in NE formation when not bound to LEM-2. Furthermore, the incidence of oocyte pronuclear collapse and embryonic lethality was significantly reduced in the *lem-2 11WH; baf-1(L58R)* mutants as compared to *chmp-711; baf-1(L58R)* mutants (Fig. 6E, F). Together, these data reveal that LEM-2-independent pools of CHMP-7 contribute to NE formation and suggest that a nucleoplasmic pool of CHMP-7 may be functional in nuclear assembly in *C. elegans*.

## Discussion

We demonstrate that BAF-1 binding to LEM-domain proteins determines the position and timing of LEM-2/EMR-1 and CHMP-7 assembly at the ‘sealing plaque,’ which forms the core domain of the NE adjacent to spindle MTs in fertilized *C. elegans* oocytes. We show that NE hole closure around spindle MTs does not depend on LEM-2, EMR-1 or CHMP-7 on their own, but BAF binding to LEM-2 and EMR-1 functions redundantly to enable hole closure. Our genetic analysis also revealed unique functions for LEM-2 and EMR-1 aside from binding to BAF-1 in NE stability and germline development. We demonstrate that CHMP-7 becomes essential to formation of the NE permeability barrier and embryonic viability in the absence of BAF-LEM domain binding. Both LEM-2 dependent and independent pools of CHMP-7 contribute to this essential function. Thus, multiple redundant mechanisms exist to prevent failure in post-meiotic NE assembly, which is essential for early embryo development.

We propose that BAF-mediated recruitment of LEM proteins and associated membranes resolves large gaps in the core domain of the NE that is obstructed by spindle microtubules while CHMP-7 functions with LEM-2 to stabilize the NE against failure in hole closure (Fig. S6D, top panel). Our prior work showed that limiting membrane biogenesis ensures successful post-meiotic hole closure (Penfield et al., 2020; Barger et al., 2022), providing further evidence that membrane feeding to narrow and close holes may be regulated to establish the NE permeability barrier. The function for BAF-LEM binding in post-meiotic hole closure is similar to its role in repair of interphase ruptures that do not contain MTs but are also micron-scale sized (Halfmann et al., 2019; Young et al., 2020). *In vitro* cross-linked BAF can exclude large macromolecules (Samwer et al., 2017) and it’s possible that BAF-LEM interactions further serve to plug large NE holes to promote sealing. Our data reveals an additional requirement for LEM-2/CHMP-7 when spindle MTs are present to stabilize the NE. Our genetic analysis also shows that different LEM-domain proteins serve distinct functions in NE assembly that are independent but partially redundant with BAF-LEM binding. How BAF-LEM interactions are further regulated by BAF phosphorylation during NE sealing and assembly requires further study.

The LEM-2 winged helix (WH) domain activates CHMP-7 to promote its ESCRT-III polymerization *in vitro* (Gatta et al., 2021; von Appen et al., 2020). However, neither LEM-2 nor CHMP-7 are required for closure of the large post-meiotic NE hole on their own (Penfield et al., 2020). LEM-2/CHMP-7 assembly may instead provide a fallback mechanism that stabilizes the NE hole should NE sealing fail (Fig. S6D). A role for CHMP-7 in restricting NE hole size has recently been suggested for NE sealing following spindle pole body extrusion in *S. pombe* (Ader et al., 2022). It is also possible that LEM-2/CHMP-7 restrict uncoordinated membrane feeding or remodel abnormal membranes at the core region that otherwise make NE assembly vulnerable to failure, especially with presence of a large hole and spindle MTs. Evidence in mammalian cells and budding yeast suggests that improper activation and mislocalization of CHMP7 can lead to harmful nuclear membrane deformations (Thaller et al., 2019; Vietri et al., 2020). Thus, the disorganized assemblies of core proteins that we observed in the absence of BAF-LEM domain binding may not only be perturbed in function for sealing, but also deleterious to NE formation. We also cannot eliminate the possibility that the BAF-L58R mutant may retain some binding to LEM domain proteins, although the accumulation of LEM domain proteins at the NE is significantly reduced. Nevertheless, our genetic evidence showing that loss of CHMP-7 exacerbates phenotypes in the BAF-L58R mutant suggest that these assemblies or other pools of CHMP-7 (see below) protect against assembly failure.

In *C. elegans,* CHMP-7 is constitutively localized in the nucleoplasm and at the INM in fully formed nuclei (Shankar et al., 2022). This localization is unlike CHMP7 in budding yeast and mammalian cells in which it is cytoplasmic or ER-associated, respectively, and thus physically segregated from LEM-2 presumably in an inactive form (Bauer et al., 2015; Gatta et al., 2021; Olmos et al., 2016; Thaller et al., 2019). CHMP7 is excluded from the nucleus in these systems through a conserved nuclear export signal and activated when it gains access to the LEM-2 WH in mitosis and upon loss of NE integrity in interphase (Gatta et al., 2021; Thaller et al., 2019; Vietri et al., 2020). The fact that CHMP-7 is constitutively nuclear-enriched and accessible to LEM-2 in *C. elegans* highlights that there may be alternative mechanisms that activate this complex upon loss of NE integrity or that cytoplasmic CHMP-7 in *C. elegans* may be specifically primed for this function. When we generated a deletion allele in *lem-2*, we discovered that LEM-2 is required for the majority of CHMP-7 to associate with the INM in *C. elegans*. Thus, LEM-2/CHMP-7 may be co-polymerized at the INM in *C. elegans* after NE sealing, but limited in the ability to recruit downstream ESCRTs.

We found that a LEM-2-WH-independent pool of CHMP-7 partially supports NE assembly (Fig. S6D, bottom panels). Prior work showed that CHMP7 binds peripherally to membranes through a hydrophobic patch that in budding yeast resembles an amphipathic helix (AH) and this association is necessary for its function (Olmos et al., 2016; Thaller et al., 2021). Prediction algorithms were unable to identify a reliable AH in a similar region for *C. elegans* CHMP-7 where a stretch of hydrophobic residues exist. Whether the nuclear or minor nuclear rim pool of CHMP-7 is the functional pool that partially supports NE assembly is unclear. Interestingly, aberrant nuclear accumulation of CHMP7 is associated with diseased neurons from patients with ALS (amyotrophic lateral sclerosis) and reduces NPC levels (Coyne et al., 2021). There may be aspects of *C. elegans* CHMP-7 that allow for it to remain inactive when soluble in the nucleus or for it to perform a unique function when unbound to membranes.

It is surprising that some oocyte-derived pronuclei assemble in the absence of both BAF-LEM domain binding and CHMP-7 even with a defective nuclear permeability barrier. This suggests that nuclear assembly is prone to failure at a stochastic rate that may depend on whether the timing of dissolution and detachment of spindle MTs is synchronized with the local stability of the NE hole. These NE irregularities sometimes prevent nuclear assembly or allow assembly of nuclei that later collapse. Our genetic experimental system of *C. elegans* allowed us to quantitively analyze NE sealing and monitor its impact on nuclear assembly and embryonic survival. Together, the redundant mechanisms that support NE assembly made evident in this study emphasize the robust nature of NE formation that is required for early development.

## Acknowledgments

We thank members of the Bahmanyar Lab for helpful discussion and critiques. We thank Victoria Puccini de Castro for help in annotating the *lem-2(tm1582)* mutant allele. We thank Sofia Sepúlveda Jacobson for help with designing the model schematic. We thank Jon Audhya and Peter Askjaer for strains and antibodies used in experiments. Some of the nematode strains were provided by the CGC, which is funded by NIH Office of Research Infrastructure Programs (P40 OD010440).

## Competing interests

The authors declare no competing or financial interests.

## Author Contributions

Conceptualization: S.R.B, S.B; Methodology: S.R.B, L.P.; Validation: S.R.B.; Formal analysis: S.R.B.; Investigation: S.R.B; Resources: S.R.B, L.P.; Data curation: S.R.B.; Writing - original draft: S.R.B, S.B; Writing – review & editing: S.R.B, L.P., S.B.; Visualization: S.R.B.; Supervision: S.B.; Project administration: S.R.B, S.B; Funding acquisition: S.R.B, S.B.

## Funding

This work was supported by an ACS Postdoctoral Fellowship (PF-21-187-01-CCB) to S.R.B and an NSF Career Award (1846010) and NIH R01 to S.B. (GM131004).

## Data availability

Almost all relevant data can be found within the article and its supplementary information. Further requests for data or reagents can be made to the corresponding author. CRISPR-Cas9 edited strains will be deposited in the CGC.

## Materials and Methods

### Strain maintenance and generation

The *C. elegans* strains used in this study are listed in Table S1. Strains were maintained at 20°C on nematode growth media (NGM) plates seeded with OP-50 *Escherichia coli.* The original MosSCI strain was maintained at 15°C.

#### MosSCI strains

The re-encoded *baf-1* transgene was cloned into pCFJ151 for integration into the Mos1 site on Chromosome I (MosSCI; (Frøkjaer-Jensen et al., 2008)). Mutagenesis was performed using the primers listed in Table S2. Injection mixes contained the following: *baf-1* transgene-containing plasmid (10 ng/μl), a plasmid encoding the Mos1 transposase (Pglh-2::transposase, pJL43.1, 10 ng/μl), and three plasmids encoding fluorescent co-injection markers: Pmyo-2::mCherry (pCFJ90, 1.4 ng/μl), Pmyo-3::mCherry (pCFJ104, 2.9 ng/μl), and Prab-3::mCherry (pGH8, 5.7 ng/μl). Injection mix was spun down (15,000 rpm, 15 minutes, 4°C) to remove particulates. Young adult hermaphrodites (EG8078) were injected and singled onto OP50 seeded plates for recovery. After ∼7-10 days, non-*unc* progeny lacking fluorescent markers were isolated and screened for transgene integration by PCR.

#### CRISPR-Cas9 deletion strains

The deletion for *lem-2* (W01G7.5) was generated using two CRISPR guides “crRNA”, which were chosen using IDT’s custom CRISPR guide algorithm (See Figure S1G). Individually, 1 μl of the purified crRNAs were annealed to 1 μl of trans-activating crRNA (tracrRNA) by incubating RNAs at 95°C for 5 minutes. A dpy-10 crRNA was used for co-CRISPR selection. An injection mix with the following components was set up at room temperature and incubated for 5 minutes: *lem-2* crRNA-1 (11.7 μM), *lem-2* crRNA-2 (11.7 μM), purified Cas9-NLS protein (qb3 Berkeley, 14.7 μM), *dpy-10* guide (3.7 μM). Finally, a *dpy-10* repair template (29 ng/mL) (Paix et al., 2015) was added and the mix was spun down (15,000 rpm) for 30 minutes at 4°C. The RNA-protein mix was injected into the gonads of N2 young adult worms, which were allowed to recover for three days. F1 progeny with a roller phenotype were singled out to individual plates. After three days, F1 mothers were genotyped by PCR. The deletion strain was sequenced and outcrossed six times to N2 worms before use and characterization.

#### CRISPR-Cas9 fluorescent knock-in strains

Fluorescent endogenous tagging of *baf-1, lem-2 and emr-1* was performed using a self-excising cassette (SEC) repair template (Dickinson and Goldstein, 2016; Hastie et al., 2019) (see Fig. S2E & S3A). Homology arms (500-800bp) were cloned into the SEC vectors (pDD268). Unique sgRNAs were cloned into the Cas9 guide plasmid pDD122 (Addgene, #47550). Following sequencing, injection plasmids were miniprepped using PureLink HiPure Plasmid Miniprep Kit (Thermo Fisher Scientific). Plasmids were combined at 100 ng/μl SEC plasmids and 50 ng/μl Cas9 plasmid and spun down (15,000 rpm) for 30 minutes at 4°C. Young adult N2 worms were injected and rescued to individual plates to recover for three days at 25°C. Plates were screened for roller progeny and positive plates were treated with approximately 200-300 μl of hygromycin B (20 mg/mL) (Thermo Fisher Scientific). Hydromycin-resistant roller worms were singled to individual plates to assess roller progeny. Plates that had roller progeny were heat-shocked at 34°C for 4 hours and allowed to recover at 20°C. Non-roller worms were singled to individual plates and progeny were screened for fluorescent protein integration by microscopy. Knock-in worms were sequence-confirmed and outcrossed four times to N2 worms.

### RNAi

Primers were designed using Primer3T to amplify a 200-1000 bp region of the gene of interest (see Table S2) using N2 gDNA as a template. The amplicon was then column purified and reversed transcribed using a T7 enzyme (MEGAscript, Life Technologies). The synthesized RNAs were purified using phenol-chloroform and resuspended in 1X soaking buffer (32.7 mM Na_2_HPO_4_, 16.5 mM KH_2_PO_4_, 6.3 mM NaCl, 14.2 mM NH_4_Cl). RNA reactions were annealed at 68°C for 10 minutes followed by 37°C for 30 minutes. dsRNAs were brought to a final concentration of ∼2000 ng/μl and stored as 2-μl aliquots at –80°C. For each experiment, a fresh aliquot was diluted to ∼1000 ng/μl using 1X soaking buffer and centrifuged at 15,000 rpm for 30 minutes at 4°C. 0.35μl of the diluted dsRNA was loaded into the back of a pulled capillary needle and injected into the gut of L4 worms. Injected worms were rescued to plates seeded with OP-50 and allowed to recover prior to imaging or lethality analysis. Knockdown time for the following targets: *baf-1* (48-72 hr), *emr-1* (24 hr), *lem-4* (24 hr), *chmp-7* (28 hr).

### Lethality and brood size quantification

L4 worms were singled (uninjected) or injected with indicated dsRNA and allowed to recover for 24 hours at 20°C. Worms were then singled to individual plates for 24 hours (day 1). Worms were transferred to another plate for a final 24 hours (day 2). Worms were then killed and plates corresponding to 24-48 hours post-injection (day 1) were counted for hatched larvae and unhatched embryos. The next day the plate corresponding to 48-72 hours post-injection (day 2) were counted. The total number of embryos and larvae were combined for each time window to calculate the brood size and the 48-72 hr embryonic lethality is reported.

### Immunoblots

#### Generation of whole worm lysate

Prior to lysate generation, L4 worms were selected 24 hours prior. For each sample, a microcentrifuge tube was filled with 30 μl of M9 Buffer and volume was marked. 35 adult worms were singled on to an indented slide filled with 75 ul of M9 + 0.1% Triton-X100. Worms were then collected into the marked microcentrifuge tube and washed three times with M9 + 0.1% Triton X-100 (200 x g, 2 minutes). After the final wash, samples were brought up to a final volume of 30 μl using M9 + 0.1% Triton. Then, 10 μl of 4X Laemmli sample buffer was added and the tubes were mixed. The samples were then sonicated at 70°C for 15 minutes, followed by incubation for five minutes at 95°C. Samples were re-sonicated at 70°C for an additional 15 minutes. Worm lysates were stored at −20°C until they were run on an SDS-PAGE protein gel.

#### Gel electrophoresis

Worm lysates were loaded on a 4-20% Mini-Protean TGX Precast Gel (Bio-Rad). The gel was run at 120V for 15 minutes to fully collapse samples and then 180V. Transfer to PDVF (Thermo Scientific) was performed at 4°C at 350mA for 1.25 hr. Membranes were blocked in 5% non-fat milk/TBST for one hour at room temperature and incubated overnight at 4°C with the following primary antibodies diluted in blocking reagent: 1 μg/mL mouse α-alpha-tubulin (DM1A, EMD Millipore), 1 μg/mL rabbit α-LEM-2 (Novus Biologicals), 1 μg/mL rabbit α-CHMP-7 (Shankar et al., 2022) and rabbit α-BAF-1 (Gorjanacz et al., 2007). The following day, membranes were briefly rinsed in TBS followed by three five-minute washes in TBST. Membranes were then incubated with appropriate secondary antibodies for 1.25 hrs at room temperature. Secondary antibodies were diluted 1:10,000 for horseradish peroxidase (HRP)-conjugated goat-anti-rabbit and HRP-conjugated goat-anti-mouse (Thermo Fischer Scientific). Membranes were again briefly rinsed in TBS followed by three five-minute washes in TBST. Membranes were incubated with Clarity Max Western ECL Substrate (BIO-RAD) for five minutes before imaging (BioRad ChemiDoc MP Imaging Systems).

### Immunofluorescence

#### Slide preparation

Microscope slides (Fisher Scientific Premium Microscope Slides Superfrost) were coated with poly-L-lysine (1 μg/mL) and dried on a heat-block. Slides were then baked at 95°C for 30 minutes and used the same day.

#### Fixation and immunofluorescence

15-20 adult worms were picked into a 4 μl drop of ddH_2_O and covered with a standard 18×18mm coverslip. Embryos were pushed out of the adult worms by pressing down on the corners of the coverslip with a pipet tip. To crack the eggshell and permeabilize the embryos, slides were placed in liquid nitrogen for ∼five-minutes. Coverslips were quickly removed by razor blade to pop off the coverslip. Slides were then fixed in pre-chilled 100% methanol at −20 °C for 20 minutes. Following fixation, slides were washed two times in 1X PBS at room temperature for 10 minutes each using a coplin jar. After the second wash, samples were blocked with 1 % BSA in PBS per slide in a humid chamber for one hour at room temperature. Slides were then incubated overnight at 4°C with primary antibodies diluted in PBS (45 μl per slide; rabbit α-LMN-1 (1 μg/mL) (Penfield et al., 2018). Following primary antibody incubation, slides were washed two times in 1X PBS+ 0.2%Tween 20 (PBST) at room temperature for 10 minutes each using a coplin jar. Following the second wash, slides were incubated at room temperature for two hours in the dark with secondary antibodies, anti-rabbit Cy3/Rhodamine, 1:200; anti-mouse FITC, 1:200 (Jackson Immunoresearch), diluted in PBS. Slides were again washed two times in PBST at room temperature for 10 minutes each in the dark. Samples were stained with 1 µg/mL Hoechst (diluted from a 1 mg/mL stock in H_2_O) for 10 minutes. Slides were washed quickly once with PBS at room temperature prior to mounting. Mounting media (Molecular Probes ProLong Diamond Antifade Reagent) was added to each sample and coverslips were adhered with clear nail-polish. Slides were dried at room temperature overnight and stored at –20°C.

### Microscopy

#### Live-cell imaging

2% agarose imaging pads were made by sandwiching molten agarose (95°C) on a glass slide. Gravid adult hermaphrodites were dissected using G10 beveled needles in 7 μl of Egg Salts (88.5 mM NaCl, 30mM KCl, 2.55 mM MgCl_2_, 2.55 mM CaCl_2_, 3.75 mM HEPES pH 7.4) on a glass slide. Select embryos were transferred to the imaging pad using a mouth pipette. Stage of interest embryos were delicately positioned using an eyelash tool and a glass coverslip was gently added on top of the imaging pad. Imaging was performed on an inverted Nikon (Melville, NY) Ti microscope with a 60X (1.4 NA) Plan Apo objective lens, a confocal scanner unit (CSU-XI, Yokogawa) with solid state 150-mW 488-nm and 100-mW 560-nm lasers, and an ORCA R-3 Digital CCD Camera (Hamamatsu). For most experiments, images were acquired every 20 seconds at five 2 μm-step z-slices. Imaging was performed in a temperature-controlled room at 20°C.

#### Live-cell meiosis imaging

Early embryo imaging, prior to eggshell formation, has been previously described (Maddox and Maddox, 2012). In brief, a circle of vasoline is drawn on a 24 x 50mm coverslip (Fisherbrand). 3 μl of Egg Salts are added to the center of the circle to dissect embryos from gravid adults. A second long coverslip is placed on top of first coverslip to form a droplet. Embryos are imaged in the suspended droplet with the vasoline preventing any harmful compression.

#### Fixed imaging

Immunofluorescent, fixed embryo imaging was conducted on an inverted Nikon Ti2 Eclipse microscope equipped with solid state 405, 445, 488, 515, 594, 561, 594, and 640 nm lasers, a Yokogawa CSU-W1 confocal scanner unit, a 60X (1.4 NA) Plan Apo objective lens, and a Prime BSI sCMOS camera (Photometrics).

### Image analysis

#### Nuclear import analysis

To determine the fluorescence intensity of NLS-LacI:GFP inside the nucleus of one-cell stage embryos, the chromatin was traced with either the freehand or circle tool in ImageJ. Camera background was determined by drawing a 50×50 pixel box in vacant areas of the time lapse. Average cytoplasmic values were determined by drawing a 20×20 pixel box inside the embryo, away from the growing nucleus. The nuclear to cytoplasmic ratio (N:C) was determined by subtracting the average camera background from each value and then the nuclear value was divided by the cytoplasmic value. To account for differences in nuclear size, the ratio was then multiplied by the nuclear area. The import of the NLS-LacI:GFP in to the pronuclei was graphed relative to pseudocleavage regression.

#### Line scan analysis of proteins at nuclear envelope and nucleoplasm

A five-pixel wide line was drawn and across the entire nucleus to determine the fluorescence intensity. Line scans (14 μm) along the maternal pronucleus were done at 200 seconds prior to pseudocleavage regression. Lines were drawn to avoid internal membranes to gather isolated nucleoplasmic values. The same line scan was used to acquire the average intensity for camera background and average was subtracted from all values. These values were then plotted against the relative position along the line. The two maximum peaks of the nuclear envelope values were averaged for the “nuclear envelope” value. ∼15-25 values were averaged in the nucleoplasm area to represent the nucleoplasmic value. The averaged nuclear envelope value was divided by the nucleoplasmic value to calculate the nuclear envelope : nucleoplasmic ratio.

#### Line scan analysis of nuclear envelope protein dynamics during meiosis and mitosis

A three-pixel wide by five-micron long line was drawn on the reforming nuclear envelope. Anaphase II onset was defined as the start of anaphase B with visible separation of chromatin away from the second polar body at the cortex. Line scans of nascent membrane at anaphase onset were drawn along chromatin mass using the line or free-hand tool in ImageJ. The same line scan was used to acquire the average intensity for camera background and the average was subtracted from all values. These values were then plotted against the relative position along the line. Line scans of sealing plaque/puncta enrichment were drawn through the enrichment to the opposite nuclear membrane using the line tool. The maximum peak values of the sealing plaque were divided by the values at the opposing nuclear rim to calculate the “LEM-2 puncta accumulation”. A similar technique was used to track sealing plaque proteins during mitosis. The free-hand line tool was used to draw a three-pixel wide line over one face of nuclear membrane alongside chromatin over time. These values were background-subtracted using an averaged camera background value. 5 values at each end of the line, representing the “non-core” regions, were averaged. The maximum intensity values along the line were divided by the non-core average to track core-domain enrichment over time (see Fig. S3E).

### Statistical Analysis

All statistical tests were performed using GraphPad Prism 9. Statistical analysis was performed on datasets with multiple samples and from independent biological repeats. Statistical tests used, sample sizes, definitions of replicates (N, n), and p values (p<0.05 as significance cutoff) are reported in figures, and/or figure legends, or text.

---

**Figure S1.**
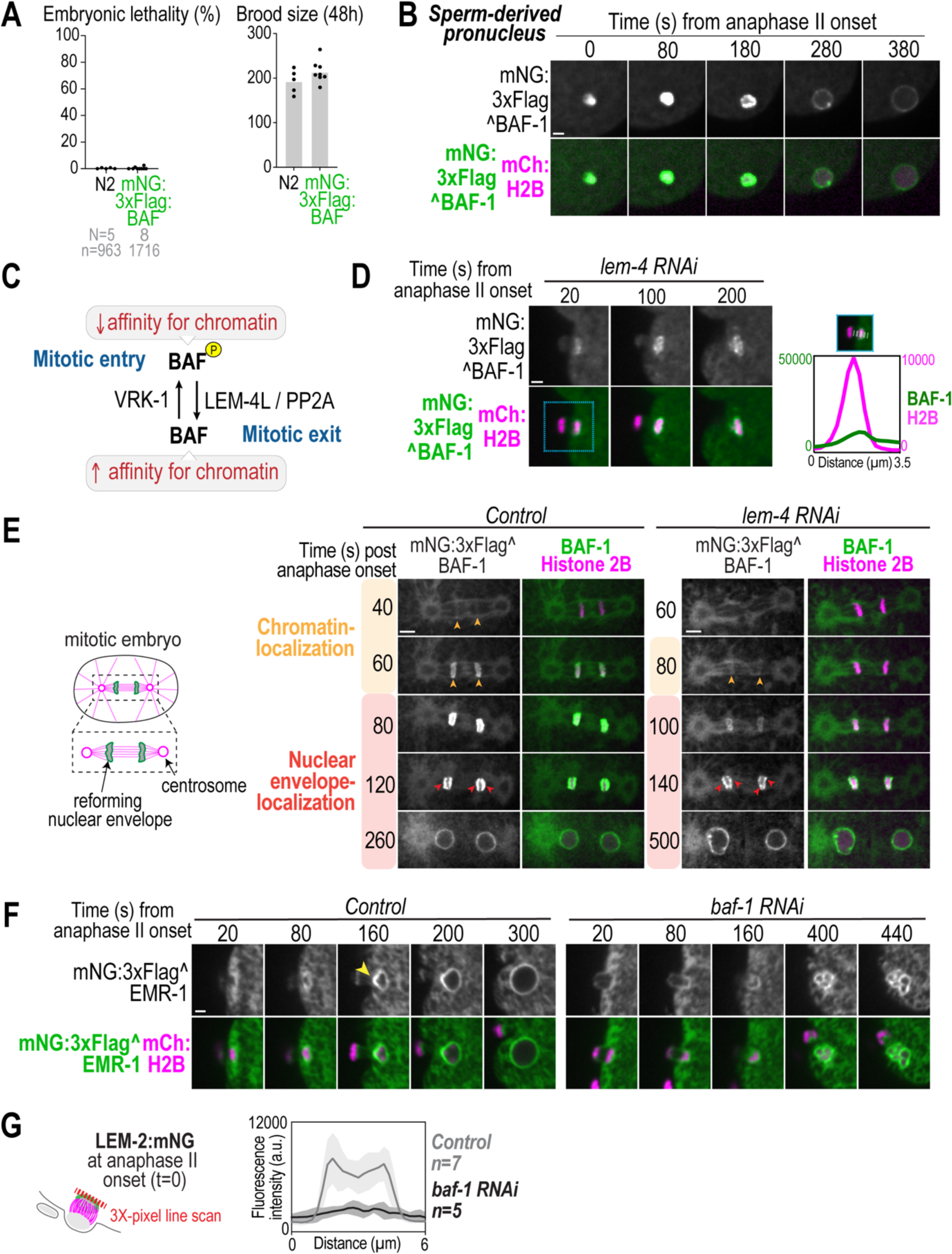
Regulation of BAF-1 dynamics, related to Figure 1. (A) Plot representing percentage of embryonic lethality and brood size in indicated conditions. N = # of worms, n = # of embryos. (B) Spinning disk confocal images from time lapse series of endogenous mNG^BAF-1 localization on sperm-derived pronucleus. Time in seconds relative to anaphase II onset. (C) Schematic of phosphoregulation of BAF-1 and the effect on chromatin interaction. (D) Left, spinning disk confocal time lapse images of mNG^BAF-1 and mCh:Histone(H)2B with *lem-4* RNAi in oocyte meiosis II. Time is in seconds relative to anaphase II. Right, plot of background-corrected line scan of indicated markers. (E) Left, schematic representation of reforming nuclear envelopes after first mitotic division in *C. elegans*. Right, spinning disk confocal time lapse series of mNG^BAF-1 and mCh:Histone(H)2B in indicated conditions. Color of arrowheads correspond to colored labels. Time in seconds relative to mitotic anaphase onset. Scale bar, 5 μm. (F) Spinning disk confocal images from time lapse series of mNG^EMR-1 dynamics during oocyte-derived pronuclear formation in indicated conditions. Yellow arrow marks sealing plaque. Time in seconds relative to anaphase II onset. Scale bar, 2 μm. (G) Left, schematic of (Right) line scan analysis (average ± SD) of LEM-2^mNG at nuclear rim at anaphase II onset in indicated conditions. n = # of embryos.

**Figure S2.**
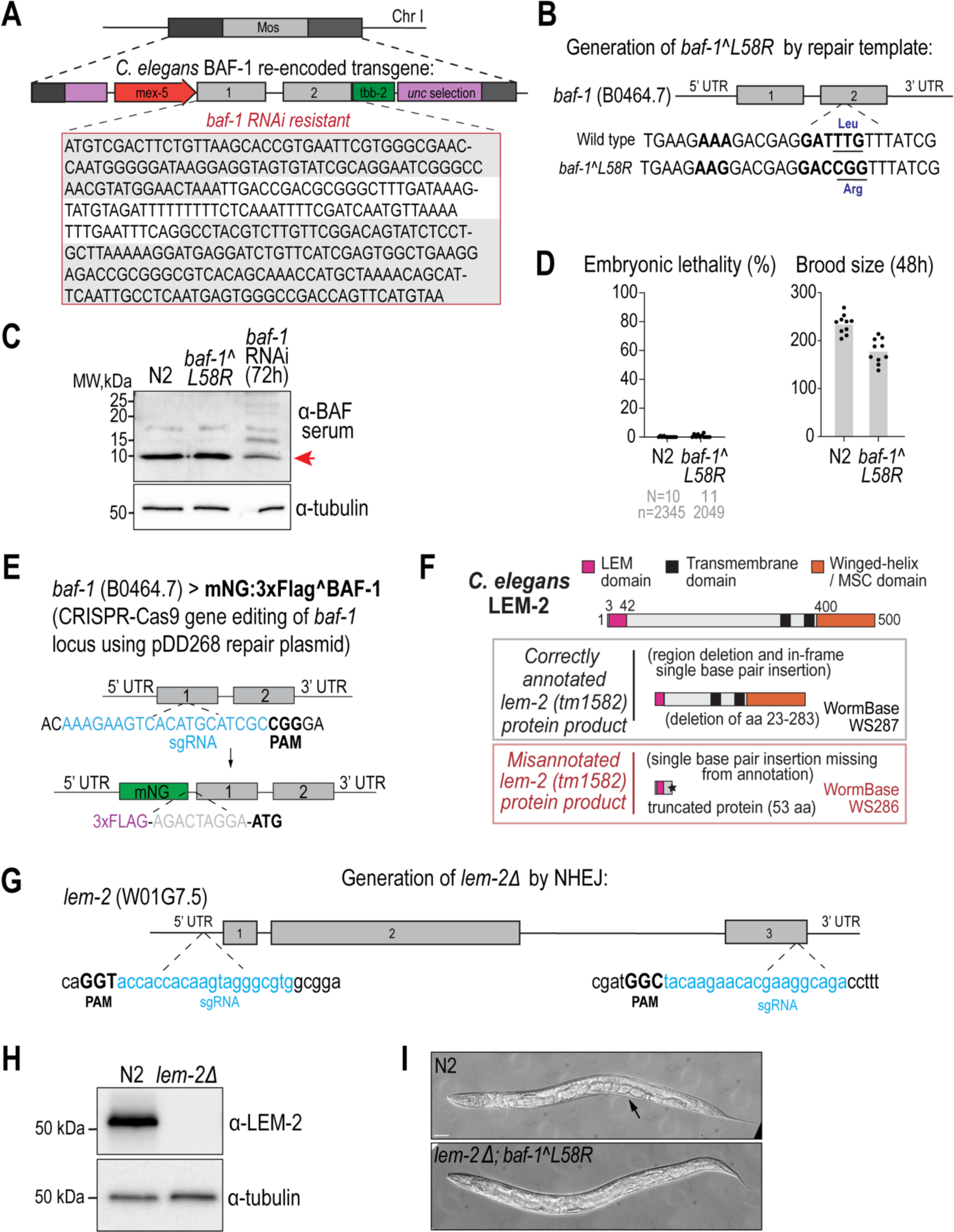
Generation and analysis of *baf-1* and *lem-2* mutant alleles. (A) Schematic showing single copy insertion of the RNAi-resistant *baf-1* transgene integrated in the MosI transposon site of chromosome I. (B) Schematic of CRISPR-Cas9 edits at endogenous *baf-1* locus to generate the *baf-1(L58R)* mutant allele. (C) Representative immunoblot of whole worm lysates incubated with antibodies against BAF in indicated conditions. N = 3 independent experiments. (D) Plots (average + replicates) representing embryonic lethality and brood size in indicated conditions. N = # of worms, n = # of embryos. (E) Schematic representation of endogenous *baf-1* locus with CRISPR guides used to insert mNG. (F) Schematic representation of LEM-2 protein domain structure. Gray box, LEM-2 mutant protein produced from correctly annotated *tm1582* mutant allele. Red box, LEM-2 mutant protein predicted from misannotated *tm1582* mutant allele. (G) Schematic representation of endogenous *lem-2* locus and CRISPR guides to excise the *lem-2* gene to generate the null allele used in this study. (H) Immunoblot of whole worm lysates incubated with antibodies made against N-terminus (aa 1-100) of LEM-2 protein (Novus Biologicals) in indicated strains. (I) Representative brightfield image of whole worms in indicated strain. Black arrow points to germline/embryos. Scale bar, 50 μm.

**Figure S3.**
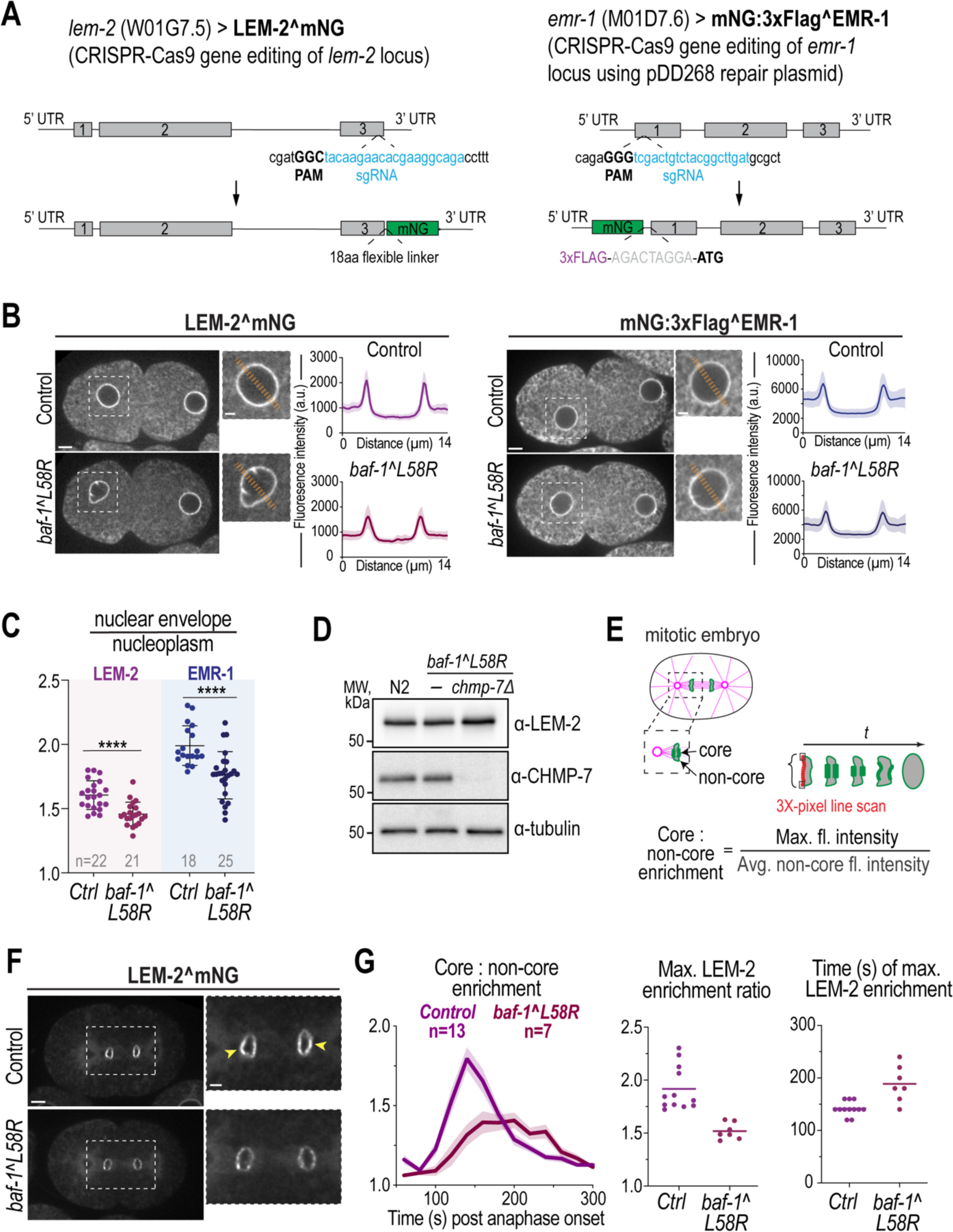
Generation of endogenous mNG tagged strains and analysis of dynamics and protein levels of EMR-1 and LEM-2 in *baf-1(L58R)* mutant strain, related to Figure 3. (A) Schematic representations of endogenous gene loci of *emr-1* and *lem-1* with CRISPR guides used to insert mNG-tag. (B) Left, representative confocal images of LEM-2^mNG and mNG^EMR-1 at oocyte- and sperm-derived pronuclei at 200 sec prior to pseudocleavage (PC) regression. Right, line scan analysis (average + SD) of LEM-2^mNG and mNG^EMR-1 at oocyte pronucleus at −200 sec releative to PC regression. n = # of embryos. (C) Plot represents average ± SD + replicates of NE:nucleoplasmic ratios of indicated proteins at oocyte pronucleus in indicated conditions. Statistical significance determined by unpaired Student’s t-test (****=p<0.0001). n = # of embryos. (D) Representative immunoblot of whole worm lysates incubated with indicated antibodies in indicated strains. N = 3 independent experiments. (E) Schematic representations of core and non-core regions on reforming NE in mitotic *C. elegans* embryo and line scan analysis of LEM-2^mNG protein dynamics shown in (G). (F) Spinning disk confocal images of endogenous LEM-2^mNG enrichment at 140 sec post anaphase onset. Yellow arrows, core enrichment. Scale bars, 5 μm. (G) Left, Plot representing average + SEM of ratio of maximum fluorescence intensity and non-core fluorescence signal measured in 20 sec intervals from anaphase onset (t = 0) in indicated conditions, Middle, Plot representing average + replicates of maximum normalized LEM:2^mNG fluorescence signal from plot on left in indicated conditions, Right, Plot representing time of maximum fluorescence signal in indicated conditions. n = # of embryos.

**Figure S4.**
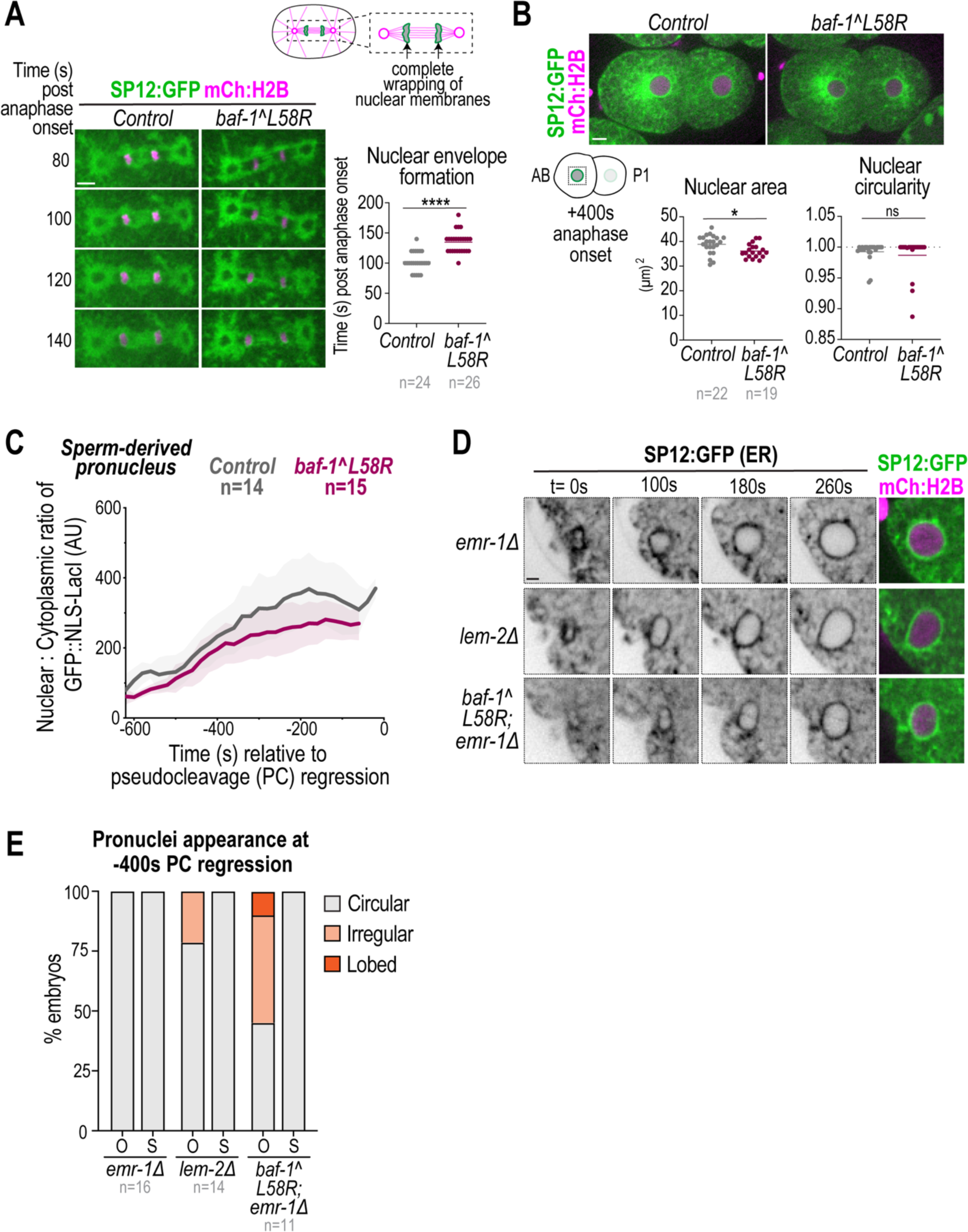
Analysis of pronuclear assembly in *emr-1Δ* and *lem-2Δ* mutants, related to Figure 4. (A) Left, representative confocal images from time-lapse series of SP12:GFP (ER marker) and mCh:Histone 2B (DNA marker) in mitotic embryos in indicated conditions. Time in seconds relative to anaphase onset. Right, plot representing average + replicates of time of complete wrapping of ER marker (SP12:GFP) around segregated chromosomes in indicated conditions. Statistical significance determined by unpaired Student’s t-test (**** = p<0.0001). n = # of embryos. (B) Above, spinning disk confocal images of indicated markers at 400 sec post anaphase onset in indicated conditions. Below, Schematic representation of 2-cell embryo and plots representing average + replicates of AB cell nuclear area and nuclear circularity at 400 sec post anaphase onset in indicated conditions. Statistical significance determined by unpaired Student’s t-test (* = p = 0.0149). ns, not significant. n = # of embryos. Scale bar, 5 μm. (C) Plot representing average ± SD of normalized nuclear GFP::NLS-LacI fluorescence in sperm-derived pronucleus for indicated conditions. Time in seconds relative to pseudocleavage (PC) regression. n = # of embryos. (D) Spinning disk confocal images from time lapse series of indicated markers during oocyte-derived pronuclear formation in indicated conditions. Scale bar, 2 μm. (E) Plot representing percentage of oocyte-derived (O) and sperm-derived (S) pronuclei categorized as circular, irregular, or lobed at −400 sec relative to pseudocleavage (PC) regression in indicated conditions. n = # of embryos.

**Figure S5.**
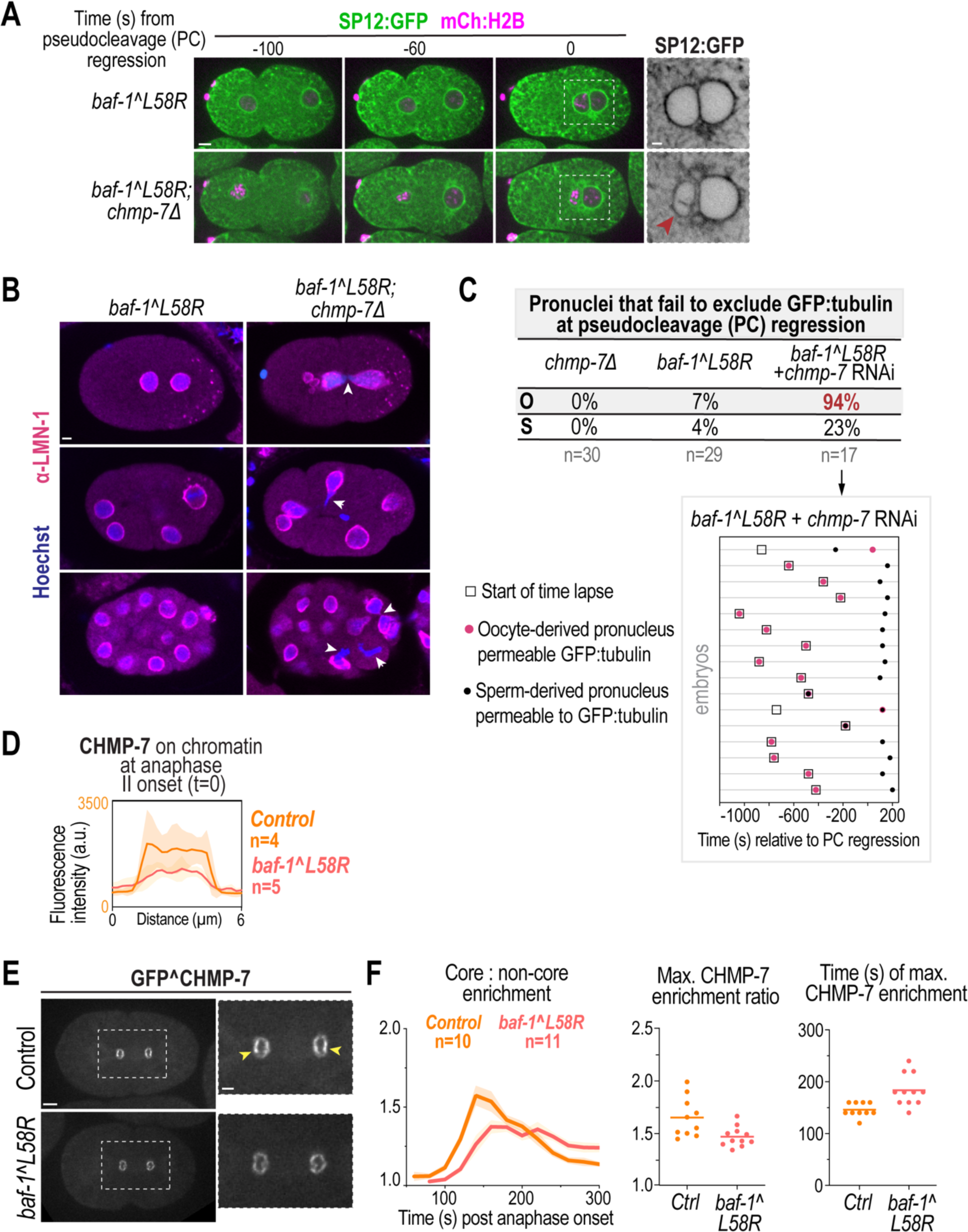
Analysis of loss of CHMP-7 and mNG^CHMP7 dynamics in meiosis and mitosis in *baf-1-L58R* mutant embryos, related to Figure 5. (A) Spinning disk confocal images from time lapse series of indicated markers in indicated conditions. Time in seconds relative to pseudocleavage (PC) regression. Scale bar, 5 μm, zoom inset image scale bar, 2 μm. (B) Confocal images of fixed 2-,8-,16+ -cell stage *C. elegans* embryos immunostained with indicated markers (LMN-1 = lamin) in indicated conditions. White arrows, chromatin bridges and impaired nuclear assembly. (C) Above, table showing percentage of oocyte-derived (O) and sperm-derived (S) pronuclei permeable to GFP:α-tubulin at pseudocleavage (PC) regression in indicated conditions. n = # of embryos. Below, Plot representing time points for individual embryos containing nuclear GFP:α-tubulin in oocyte-(red circles) or sperm-derived pronuclei (black circles) relative to start of time-lapse movie (square). Time in seconds relative to pseudocleavage (PC) regression. (D) Background-corrected line scan analysis (average ± SD) of GFP^CHMP-7 on the nuclear rim anaphase II onset (white arrows in Figure 5F) in indicated conditions. n = # of embryos. (E) Spinning disk confocal images of GFP^CHMP-7 at mitotic nuclear formation at 140 sec post anaphase onset. Yellow arrows, core domain. Scale bars, 5 μm. (F) Left, Plot representing average + SEM of ratio of maximum fluorescence intensity and non-core fluorescence signal measured in 20 sec intervals from anaphase onset (t = 0) in indicated conditions, Middle, Plot representing average + replicates of maximum normalized GFP^CHMP-7 fluorescence signal from plot on left in indicated conditions. Right, Plot representing time of maximum fluorescence signal in indicated conditions. n = # of embryos. n = # of embryos.

**Figure S6.**
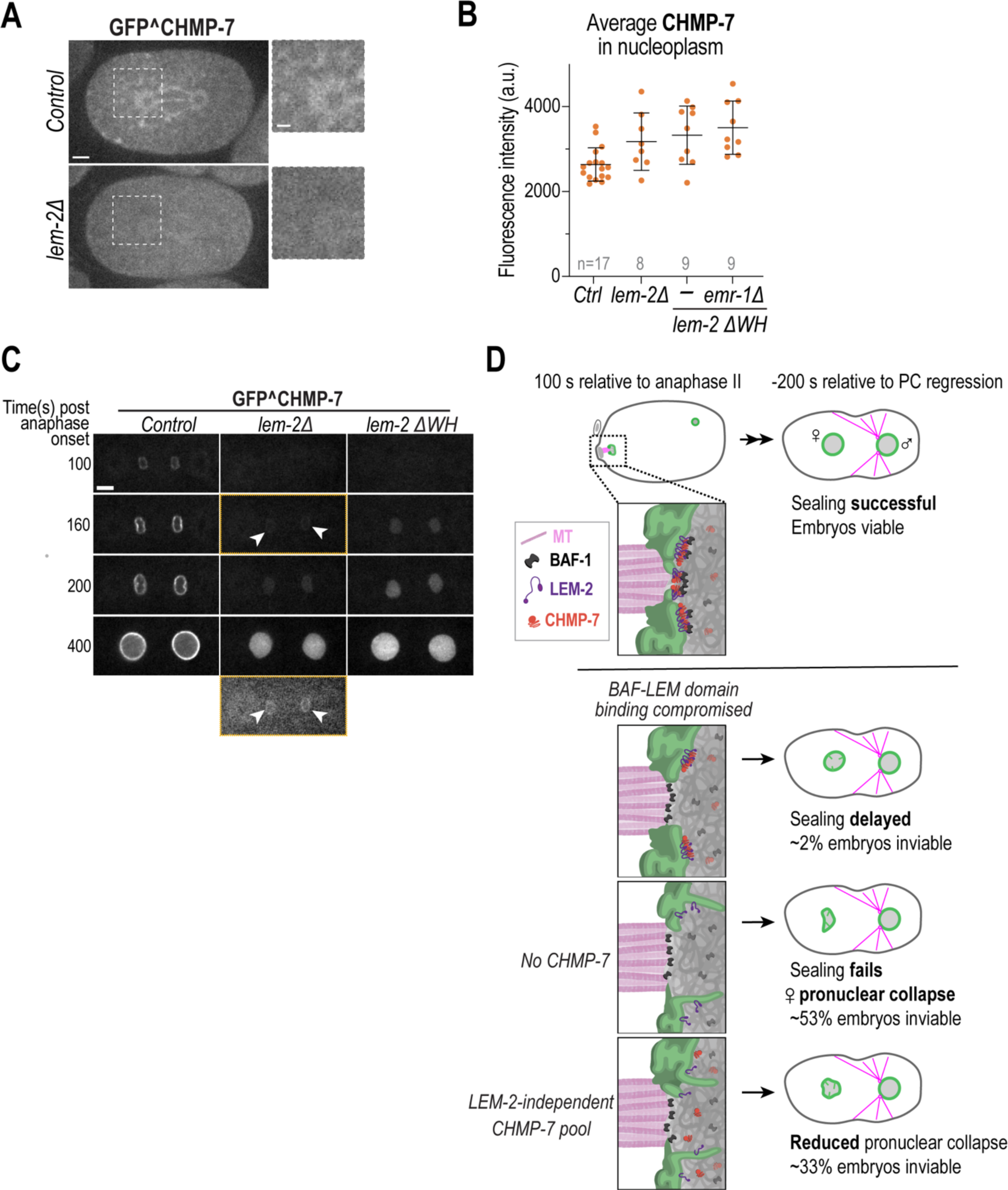
CHMP-7 localization and dependence on LEM-2, related to Figure 6. (A) Spinning disk confocal images of endogenous GFP^CHMP-7 in control and *lem-2Δ* embryos at anaphase onset in mitotic embryos. Zoom inset of spindle pole. Scale bar, 5 μm, zoom scale bar, 2 μm. (B) Plot representing average ± SD + replicates of nucleoplasmic GFP^ CHMP-7 levels in oocyte-derived pronuclei −200 sec relative to pseudocleavage (PC) regression. n = # of embryos. (C) Spinning disk confocal images from time series of GFP^CHMP-7 in indicated conditions. Time in seconds relative to anaphase onset. Scale bar, 5 μm. White arrowheads point to faint fluorescence signal of GFP^CHMP-7 prior to nuclear accumulation. Yellow outlined panel reproduced below with brightness/contrast adjusted. (D) Schematic representation of oocyte-derived pronuclear sealing in different mutant backgrounds analyzed in this study (left). Right, zoom inset of sealing plaque adjacent to spindle microtubules. LEM domain proteins (only LEM-2 is shown for simplicity) attach incoming ER membranes to BAF to narrow the nuclear envelope hole. CHMP-7/LEM-2 stabilize the nuclear envelope hole and remodel abnormal membranes, while a LEM-2 independent pool of CHMP-7 promotes nuclear stability through an unknown mechanism.

## Supplemental Movie Legends

**Movie S1. BAF-1 dynamics at oocyte- and sperm-derived pronuclei during pronuclear formation,** related to Figure 1. Spinning disk confocal fluorescence time series of mNG^BAF-1 (green) and mCherry:Histone2B (magenta) in fertilized oocyte. Time lapse shows mNG^BAF-1 enrichment on chromatin transitioning to nuclear membranes during oocyte-(left) and sperm-(right) derived pronuclear formation. Images were acquired every 20 s – playback rate is 80X real time. Scale bar, 5 μm.

**Movie S2. BAF-1 is required for LEM-2 enrichment at the sealing plaque following meiosis II,** related to Figure 1. Spinning disk confocal fluorescence time series of endogenously tagged LEM-2^mNG (green) and mCherry:Histone2B (magenta) in fertilized oocytes. Time lapse shows oocyte-derived pronuclear formation, starting at anaphase II onset, as chromatin moves away from the cortex, and ending with a sealed pronucleus in indicated conditions. LEM-2^mNG does not accumulate at a discrete focus in the reforming oocyte-derived pronucleus, which undergoes assembly failure. Images were acquired every 20 s – playback rate is 80X real time. Scale bar, 2 μm.

**Movie S3. Reduced and delayed LEM-2^mNG enrichment at the sealing plaque in *baf-1^L58R* embryos,** related to Figure 3. Spinning disk confocal fluorescence images of LEM-2^mNG (green) and mCherry:Histone2B (magenta) in fertilized oocytes in indicated conditions. Time lapse shows oocyte-derived pronuclear formation, starting at anaphase II onset. Images were acquired every 20 s – playback rate is 80X real time. Scale bar, 2 μm.

**Movie S4. Oocyte pronuclear formation in control, *baf-1^L58R*, *baf-1^L58R* + *lem-4* RNAi embryos,** related to Figure 4. Spinning disk confocal fluorescence time series of embryos expressing SP12:GFP (ER in grey, inverted) in indicated conditions. Images were acquired every 20 s – playback rate is 80X real time. Scale bar, 2 μm.

**Movie S5. Oocyte pronuclear collapse occurs in *chmp-7Δ*; *baf-1^L58R* embryos,** related to Figure 5. Spinning disk confocal fluorescence time series of SP12:GFP (ER, green) and mCherry:Histone2B (magenta) marking oocyte-derived pronuclei in indicated conditions. Time lapse shows migration of oocyte- and sperm-derived pronuclei ending in pronuclear meeting approximately at PC regression. Images were acquired every 20 s – playback rate is 80X real time. Scale bar, 5 μm.

**Movie S6. Reduced and delayed CHMP-7 enrichment at the sealing plaque in *baf-1^L58R* embryos,** related to Figure 5. Spinning disk confocal fluorescence time series of GFP^CHMP-7 (green) and mCherry:Histone2B (magenta) after anaphase II onset in fertilized oocytes in indicated conditions. Time lapse shows oocyte-derived pronuclear formation, starting at anaphase II onset as chromatin moves away from the cortex and ending with a formed pronucleus in indicated conditions. Images were acquired every 20s – playback rate is 80X real time. Scale bar, 2 μm.

**Movie S7. Reliance of GFP^CHMP-7 on the Winged Helix domain of LEM-2 to localize and enrich at the sealing plaque following meiosis II,** related to Figure 6. Spinning disk confocal fluorescence time series of GFP^CHMP-7 (green) and mCherry:Histone2B (magenta) after anaphase II onset in fertilized oocytes in indicated conditions. Left, gray scale images of GFP^CHMP-7, Right, merged. Time lapse shows oocyte-derived pronuclear formation, starting at anaphase II onset as chromatin moves away from the cortex and ending with a formed pronucleus in *lem-2 ΔWH* mutant. Left, GFP^CHMP-7 (grey), Right, merged. Images were acquired every 20 s – playback rate is 80X real time. Scale bar, 2 μm.

---

**Table S1.**
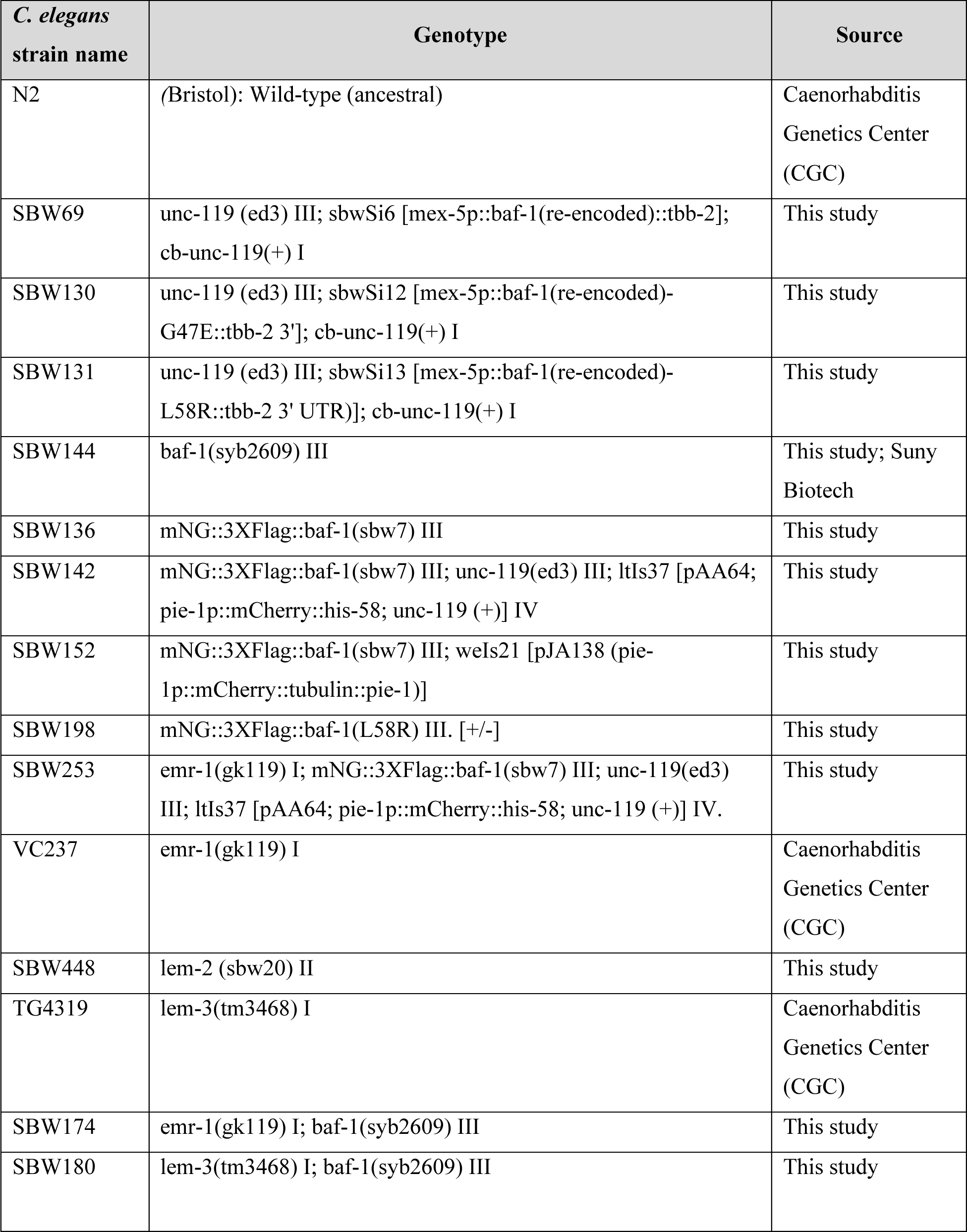
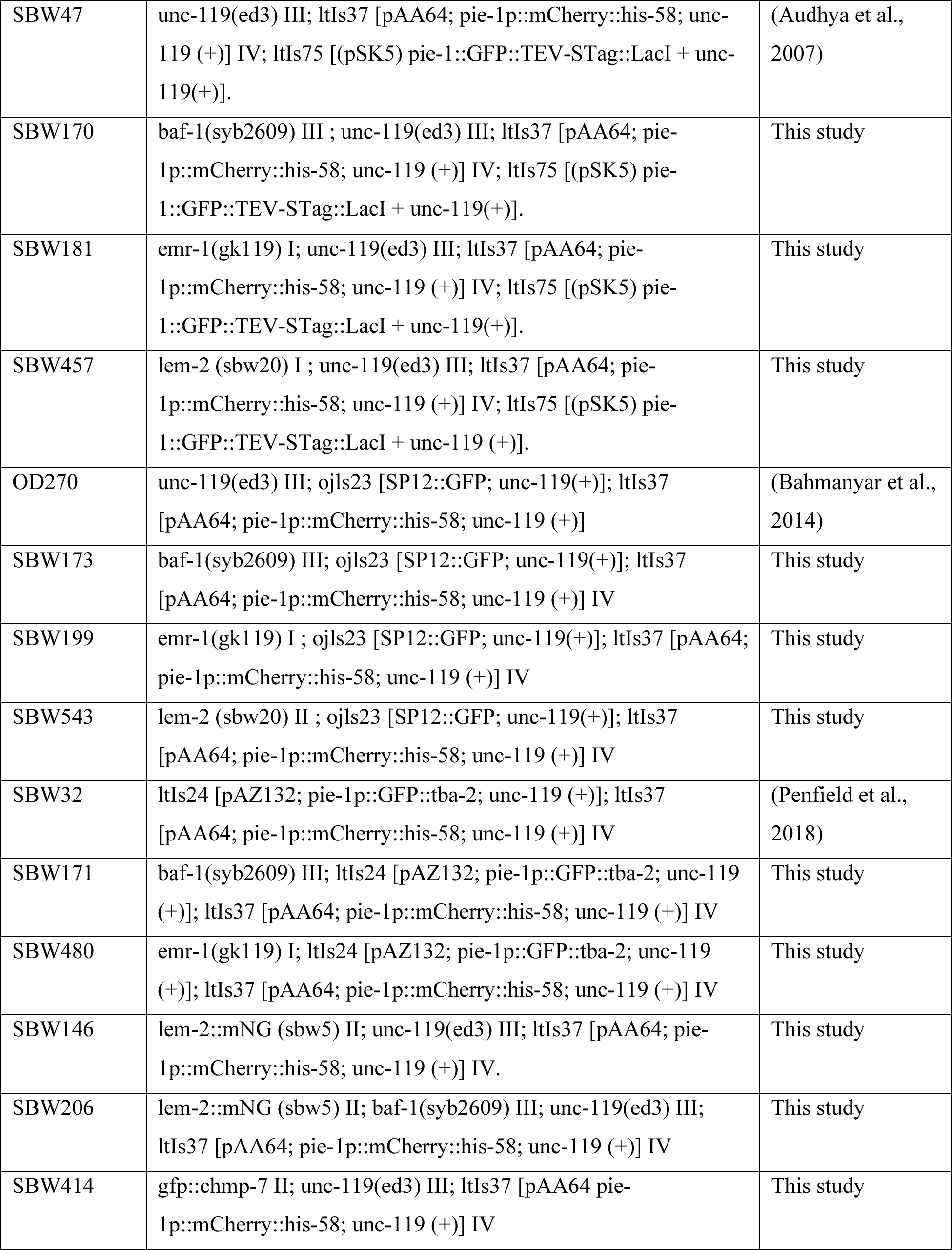
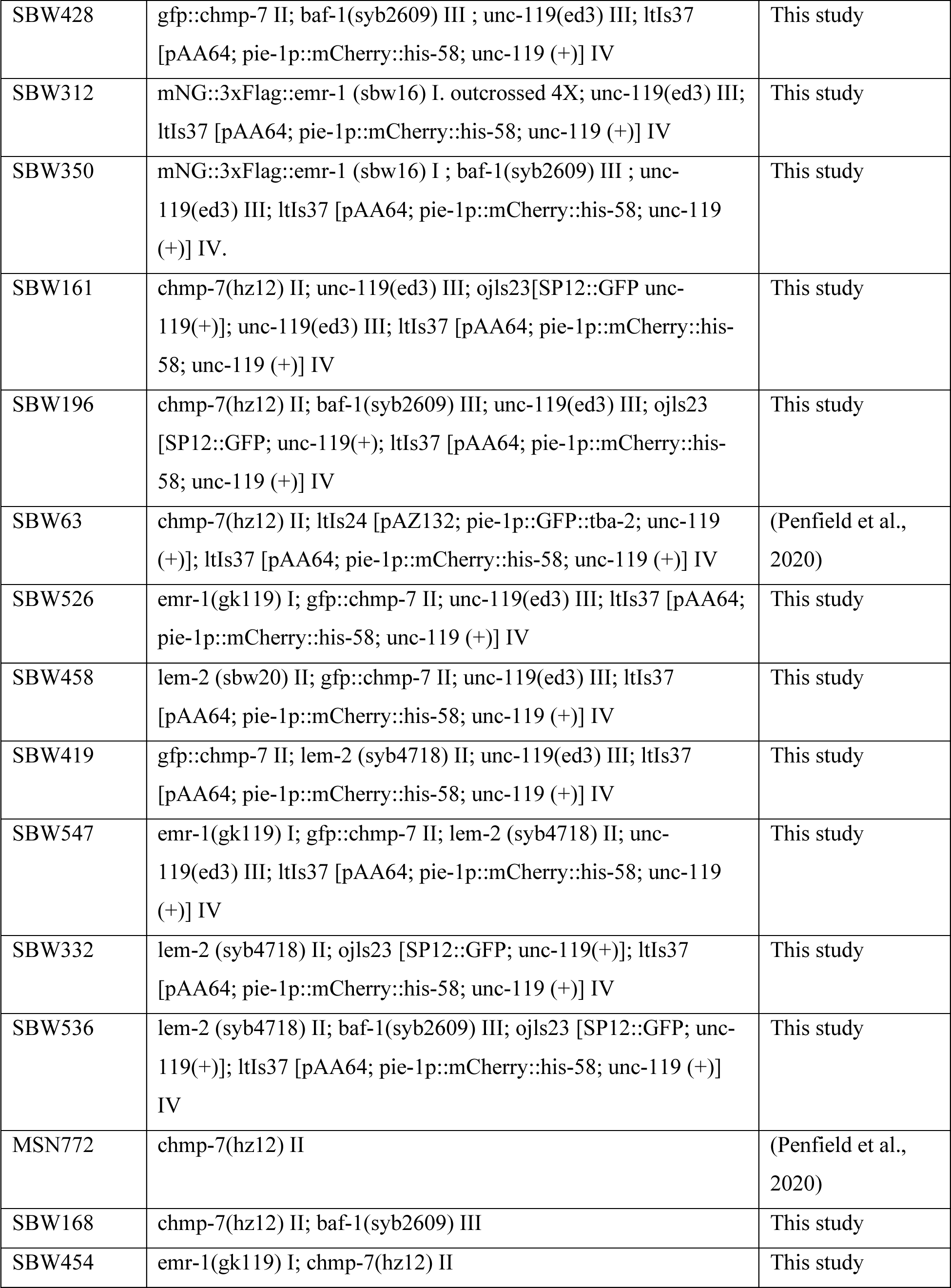
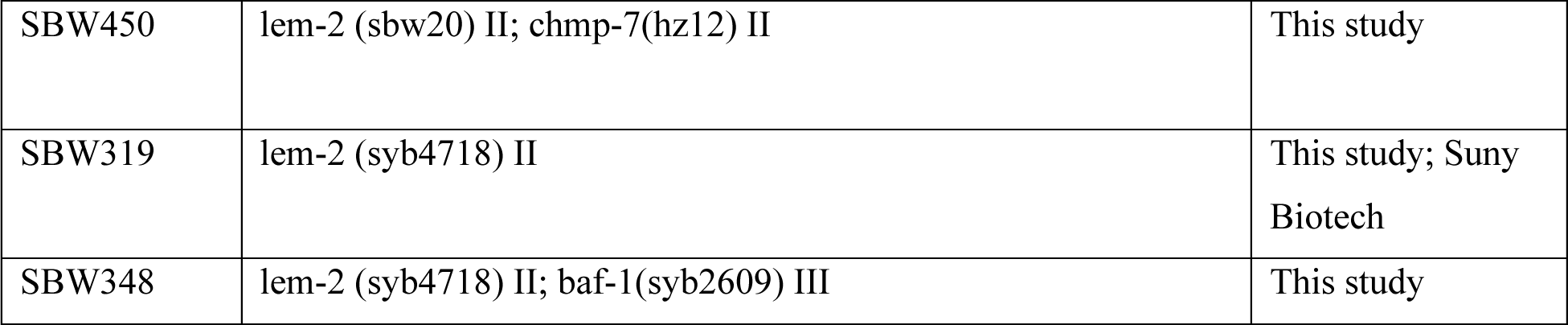
*C. elegans* strains used in this study.

| *C. elegans*<br>strain name | Genotype | Source |
| --- | --- | --- |
| N2 | (Bristol): Wild-type (ancestral) | Caenorhabditis<br>Genetics Center<br>(CGC) |
| SBW69 | unc-119 (ed3) III; sbwSi6 [mex-5p::baf-1(re-encoded)::tbb-2];<br>cb-unc-119(+) I | This study |
| SBW130 | unc-119 (ed3) III; sbwSi12 [mex-5p::baf-1(re-encoded)-<br>G47E::tbb-2 3']; cb-unc-119(+) I | This study |
| SBW131 | unc-119 (ed3) III; sbwSi13 [mex-5p::baf-1(re-encoded)-<br>L58R::tbb-2 3' UTR)]; cb-unc-119(+) I | This study |
| SBW144 | baf-1(syb2609) III | This study; Suny<br>Biotech |
| SBW136 | mNG::3XFlag::baf-1(sbw7) III | This study |
| SBW142 | mNG::3XFlag::baf-1(sbw7) III; unc-119(ed3) III; ltIs37 [pAA64;<br>pie-1p::mCherry::his-58; unc-119 (+)] IV | This study |
| SBW152 | mNG::3XFlag::baf-1(sbw7) III; weIs21 [pJA138 (pie-1p::mCherry::tubulin::pie-1)] | This study |
| SBW198 | mNG::3XFlag::baf-1(L58R) III. [+/-] | This study |
| SBW253 | emr-1(gk119) I; mNG::3XFlag::baf-1(sbw7) III; unc-119(ed3)<br>III; ltIs37 [pAA64; pie-1p::mCherry::his-58; unc-119 (+)] IV. | This study |
| VC237 | emr-1(gk119) I | Caenorhabditis<br>Genetics Center<br>(CGC) |
| SBW448 | lem-2 (sbw20) II | This study |
| TG4319 | lem-3(tm3468) I | Caenorhabditis<br>Genetics Center<br>(CGC) |
| SBW174 | emr-1(gk119) I; baf-1(syb2609) III | This study |
| SBW180 | lem-3(tm3468) I; baf-1(syb2609) III | This study |
| SBW47 | unc-119(ed3) III; ltIs37 [pAA64; pie-1p::mCherry::his-58; unc-119 (+)] IV; ltIs75 [(pSK5) pie-1::GFP::TEV-STag::LacI + unc-119(+)]. | (Audhya et al., 2007) |
| SBW170 | baf-1(syb2609) III ; unc-119(ed3) III; ltIs37 [pAA64; pie-1p::mCherry::his-58; unc-119 (+)] IV; ltIs75 [(pSK5) pie-1::GFP::TEV-STag::LacI + unc-119(+)]. | This study |
| SBW181 | emr-1(gk119) I; unc-119(ed3) III; ltIs37 [pAA64; pie-1p::mCherry::his-58; unc-119 (+)] IV; ltIs75 [(pSK5) pie-1::GFP::TEV-STag::LacI + unc-119(+)]. | This study |
| SBW457 | lem-2 (sbw20) I ; unc-119(ed3) III; ltIs37 [pAA64; pie-1p::mCherry::his-58; unc-119 (+)] IV; ltIs75 [(pSK5) pie-1::GFP::TEV-STag::LacI + unc-119 (+)]. | This study |
| OD270 | unc-119(ed3) III; ojls23 [SP12::GFP; unc-119(+)]; ltIs37 [pAA64; pie-1p::mCherry::his-58; unc-119 (+)] | (Bahmanyar et al., 2014) |
| SBW173 | baf-1(syb2609) III; ojls23 [SP12::GFP; unc-119(+)]; ltIs37 [pAA64; pie-1p::mCherry::his-58; unc-119 (+)] IV | This study |
| SBW199 | emr-1(gk119) I ; ojls23 [SP12::GFP; unc-119(+)]; ltIs37 [pAA64; pie-1p::mCherry::his-58; unc-119 (+)] IV | This study |
| SBW543 | lem-2 (sbw20) II ; ojls23 [SP12::GFP; unc-119(+)]; ltIs37 [pAA64; pie-1p::mCherry::his-58; unc-119 (+)] IV | This study |
| SBW32 | ltIs24 [pAZ132; pie-1p::GFP::tba-2; unc-119 (+)]; ltIs37 [pAA64; pie-1p::mCherry::his-58; unc-119 (+)] IV | (Penfield et al., 2018) |
| SBW171 | baf-1(syb2609) III; ltIs24 [pAZ132; pie-1p::GFP::tba-2; unc-119 (+)]; ltIs37 [pAA64; pie-1p::mCherry::his-58; unc-119 (+)] IV | This study |
| SBW480 | emr-1(gk119) I; ltIs24 [pAZ132; pie-1p::GFP::tba-2; unc-119 (+)]; ltIs37 [pAA64; pie-1p::mCherry::his-58; unc-119 (+)] IV | This study |
| SBW146 | lem-2::mNG (sbw5) II; unc-119(ed3) III; ltIs37 [pAA64; pie-1p::mCherry::his-58; unc-119 (+)] IV. | This study |
| SBW206 | lem-2::mNG (sbw5) II; baf-1(syb2609) III; unc-119(ed3) III; ltIs37 [pAA64; pie-1p::mCherry::his-58; unc-119 (+)] IV | This study |
| SBW414 | gfp::chmp-7 II; unc-119(ed3) III; ltIs37 [pAA64 pie-1p::mCherry::his-58; unc-119 (+)] IV | This study |
| SBW428 | gfp::chmp-7 II; baf-1(syb2609) III ; unc-119(ed3) III; ltIs37 [pAA64; pie-1p::mCherry::his-58; unc-119 (+)] IV | This study |
| SBW312 | mNG::3xFlag::emr-1 (sbw16) I. outcrossed 4X; unc-119(ed3) III; ltIs37 [pAA64; pie-1p::mCherry::his-58; unc-119 (+)] IV | This study |
| SBW350 | mNG::3xFlag::emr-1 (sbw16) I ; baf-1(syb2609) III ; unc-119(ed3) III; ltIs37 [pAA64; pie-1p::mCherry::his-58; unc-119 (+)] IV. | This study |
| SBW161 | chmp-7(hz12) II; unc-119(ed3) III; ojls23[SP12::GFP unc-119(+)]; unc-119(ed3) III; ltIs37 [pAA64; pie-1p::mCherry::his-58; unc-119 (+)] IV | This study |
| SBW196 | chmp-7(hz12) II; baf-1(syb2609) III; unc-119(ed3) III; ojls23 [SP12::GFP; unc-119(+); ltIs37 [pAA64; pie-1p::mCherry::his-58; unc-119 (+)] IV | This study |
| SBW63 | chmp-7(hz12) II; ltIs24 [pAZ132; pie-1p::GFP::tba-2; unc-119 (+)]; ltIs37 [pAA64; pie-1p::mCherry::his-58; unc-119 (+)] IV | (Penfield et al., 2020) |
| SBW526 | emr-1(gk119) I; gfp::chmp-7 II; unc-119(ed3) III; ltIs37 [pAA64; pie-1p::mCherry::his-58; unc-119 (+)] IV | This study |
| SBW458 | lem-2 (sbw20) II; gfp::chmp-7 II; unc-119(ed3) III; ltIs37 [pAA64; pie-1p::mCherry::his-58; unc-119 (+)] IV | This study |
| SBW419 | gfp::chmp-7 II; lem-2 (syb4718) II; unc-119(ed3) III; ltIs37 [pAA64; pie-1p::mCherry::his-58; unc-119 (+)] IV | This study |
| SBW547 | emr-1(gk119) I; gfp::chmp-7 II; lem-2 (syb4718) II; unc-119(ed3) III; ltIs37 [pAA64; pie-1p::mCherry::his-58; unc-119 (+)] IV | This study |
| SBW332 | lem-2 (syb4718) II; ojls23 [SP12::GFP; unc-119(+)]; ltIs37 [pAA64; pie-1p::mCherry::his-58; unc-119 (+)] IV | This study |
| SBW536 | lem-2 (syb4718) II; baf-1(syb2609) III; ojls23 [SP12::GFP; unc-119(+)]; ltIs37 [pAA64; pie-1p::mCherry::his-58; unc-119 (+)] IV | This study |
| MSN772 | chmp-7(hz12) II | (Penfield et al., 2020) |
| SBW168 | chmp-7(hz12) II; baf-1(syb2609) III | This study |
| SBW454 | emr-1(gk119) I; chmp-7(hz12) II | This study |
| SBW450 | lem-2 (sbw20) II; chmp-7(hz12) II | This study |
| SBW319 | lem-2 (syb4718) II | This study; Suny<br>Biotech |
| SBW348 | lem-2 (syb4718) II; baf-1(syb2609) III | This study |

**Table S2.**
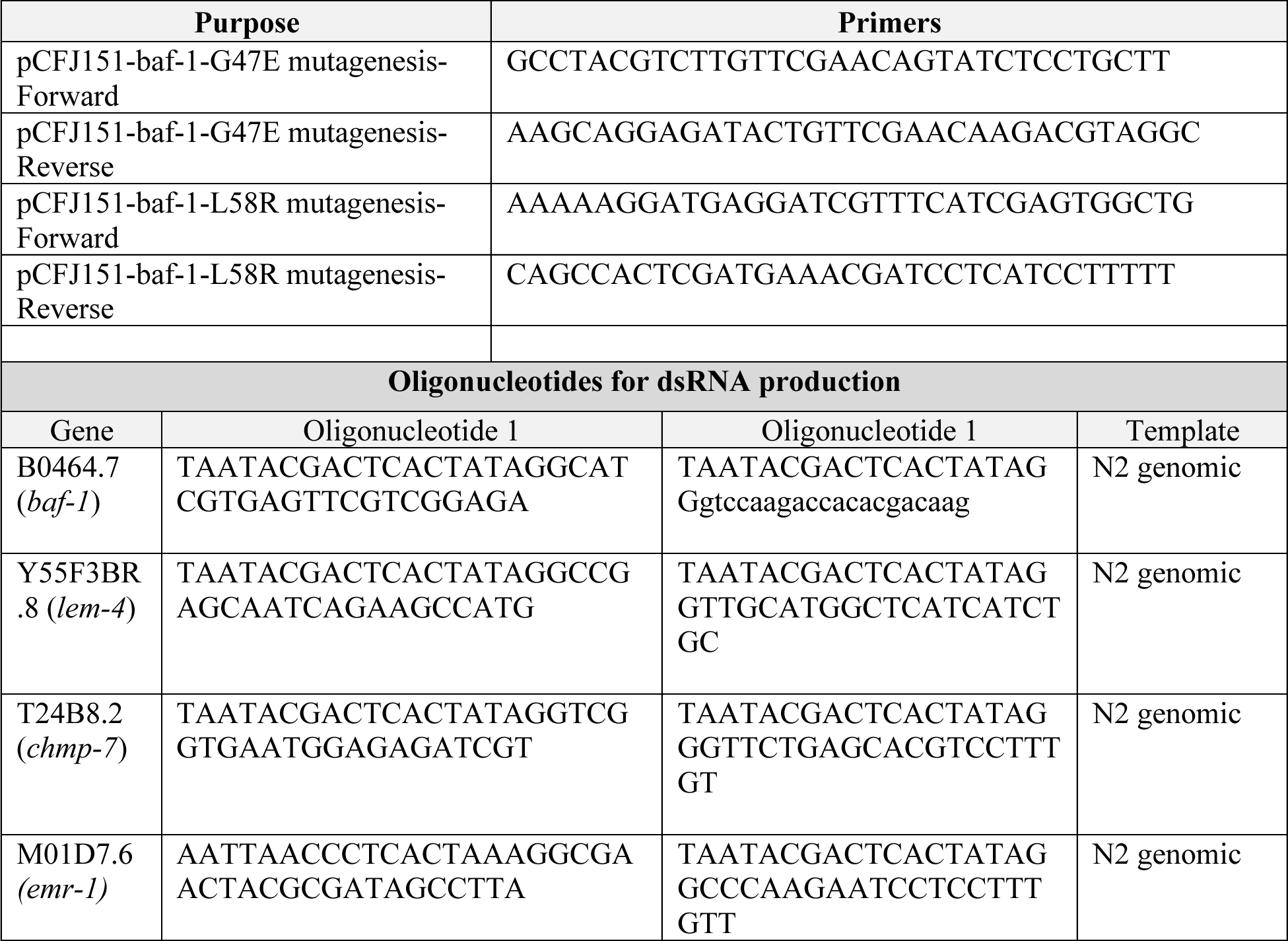
Oligonucleotides used in this study.

**Table S3.**
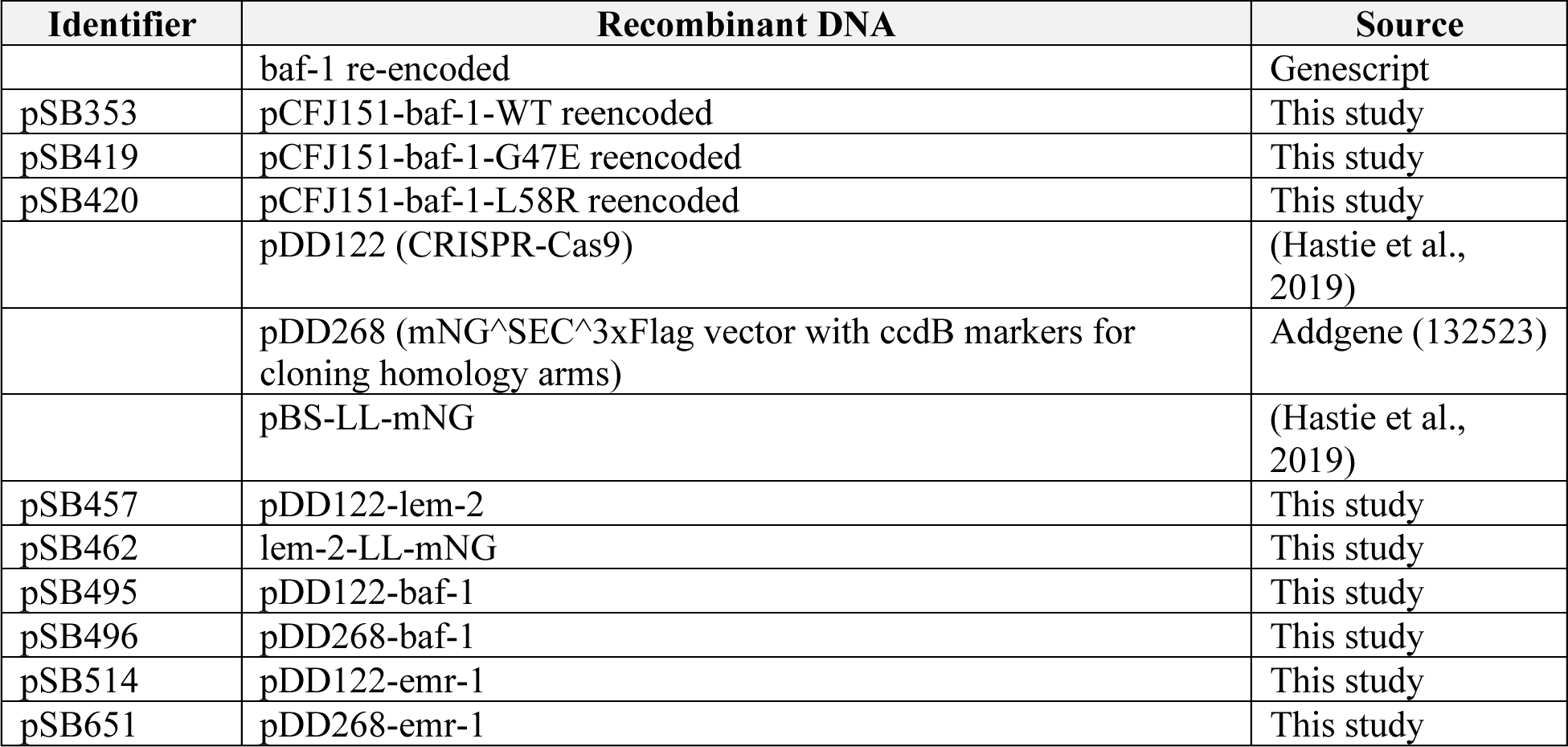
Recombinant DNA used in this study.

## Notes

### Competing Interest Statement

The authors have declared no competing interest.

